# 142 telomere-to-telomere assemblies reveal the genome structural landscape in *Saccharomyces cerevisiae*

**DOI:** 10.1101/2022.10.04.510633

**Authors:** Samuel O’Donnell, Jia-Xing Yue, Omar Abou Saada, Nicolas Agier, Claudia Caradec, Thomas Cokelaer, Matteo De Chiara, Stéphane Delmas, Fabien Dutreux, Téo Fournier, Anne Friedrich, Etienne Kornobis, Jing Li, Zepu Miao, Lorenzo Tattini, Joseph Schacherer, Gianni Liti, Gilles Fischer

**Author notes:** co-first (Samuel O’Donnell, Jia-Xing Yue). co-last, co-corresponding.

## Abstract

As population genomics is transitioning from single reference genomes to pangenomes, major improvements in terms of genome contiguity, phylogenetic sampling, haplotype phasing and structural variant (SV) calling are required. Here, we generated the *Saccharomyces cerevisiae* Reference Assembly Panel (ScRAP) comprising 142 reference-quality genomes from strains of various geographic and ecological origins that faithfully represent the genomic diversity and complexity of the species. The ca. 4,800 non-redundant SVs we identified impact the expression of genes near the breakpoints and contribute to gene repertoire evolution through disruptions, duplications, fusions and horizontal transfers. We discovered frequent cases of complex aneuploidies, preferentially involving large chromosomes that underwent large SVs. We also characterized the evolutionary dynamics of complex genomic regions that classically remain unassembled in short read-based projects, including the 5 Ty families and the 32 individual telomeres. Overall, the ScRAP represents a crucial step towards establishing a high-quality, unified and complete S. cerevisiae pangenome.

## INTRODUCTION

Third generation single-molecule long-read sequencing greatly improves genome assembly and provides access to gapless genome sequences, including repetitive and highly variable chromosomal regions that generally remain unassembled. This is best exemplified in the rapid increase in contiguity of the human genome (Chaisson *et al*., 2015), especially thanks to ultra-long reads from Oxford Nanopore Technology (ONT) (Jain *et al*., 2018). Recently, the T2T consortium released the first complete ‘telomere-to-telomere’ assembly of two human chromosomes (Logsdon *et al*., 2020, 2021; Miga *et al*., 2020), soon followed by the release of the first gapless human genome, including nearly 200 Mb of novel sequences (Nurk *et al*., 2022). Complex plant genomes have also seen improvements in assembly contiguity due to increasing read length (Choi *et al*., 2020; Qin *et al*., 2021; Rousseau-Gueutin *et al*., 2020). Classical model organisms including drosophilid species and the green algae *Chlamydomonas reinhardtii,* benefitted from ONT long-read sequencing with novel genome assemblies far more complete and contiguous than the ones that were previously available (Kim *et al*., 2021; O’Donnell *et al*., 2020).

These advancements in the ease of producing more contiguous, reference-like assemblies, has allowed few species to have multiple contiguous genomes. As expected, classical model organisms and species of anthropocentric and economic importance such as *Escherichia coli* (Wang *et al*., 2021), *Drosophila melanogaster* (Rech *et al*., 2022; Kim *et al*., 2021), *Solanum lycopersicum* (tomato) (Alonge *et al*., 2020), *Glycine max* (soybean) (Liu *et al*., 2020), *Oryza sativa* (rice) (Qin *et al*., 2021; Zhang *et al*., 2022), *Bombyx mori* (silkworm) (Tong *et al*., 2022) and humans (Audano *et al*., 2019; Beyter *et al*., 2021; Wong *et al*., 2018) already have several contiguous genomes within each species. As has often been the case in genetics and genomics, the baker’s yeast, *Saccharomyces cerevisiae*, is at the forefront of the field currently totaling 68 long-read genome assemblies of non-reference strains (Abou Saada *et al*., 2021; Bendixsen *et al*., 2021; Berlin *et al*., 2015; Czaja *et al*., 2020; Istace *et al*., 2017; Jenjaroenpun *et al*., 2018; Lee *et al*., 2022; Shao *et al*., 2018; Yue *et al*., 2017; Zhang and Emerson, 2019; Heasley and Argueso, 2022). This data has been used to quantify contiguity improvements of long-read sequencing over short-read data (Istace *et al*., 2017), create genome-wide maps of transposable elements (Bendixsen *et al*., 2021; Czaja *et al*., 2020; Istace *et al*., 2017), characterize subtelomeric regions (Yue *et al*., 2017), phase haplotypes and detect large SVs (Bendixsen *et al*., 2021; Istace *et al*., 2017; Jenjaroenpun *et al*., 2018; Yue *et al*., 2017; Heasley and Argueso, 2022).

However, several limitations remain to be overcome in *S. cerevisiae* population genomics. The contiguity of the available genome assemblies varies widely (range: 16-106 scaffolds) and only a small subset of them reached a chromosome-level scaffold contiguity. The sampling remains limited in terms of phylogeny as many clades of *S. cerevisiae* lack a representative reference genome. No polyploid genomes have been included although they represented 11.5% of the isolates (Peter *et al*., 2018, 011). Last, phasing haplotypes of diploid and polyploid genomes is challenging and as a result, information about heterozygosity has mostly remained hidden in haplotype-collapsed genome assemblies. Although in its infancy, reconstructing haplotypes has already been shown to be important in the detection of large insertions and SVs (Chin *et al*., 2016; Cretu Stancu *et al*., 2017; Garg *et al*., 2021; Kitzman *et al*., 2011; Koren *et al*., 2018; Porubsky *et al*., 2021) and in the understanding of the admixture process in human (Green *et al*., 2010). In *S. cerevisiae* the utility of phasing with long-reads has been used to accurately identify the European and Asian parental origins of polyploid ale brewing strains (Fay *et al*., 2019) and to characterize interhomolog variations such as heterozygous Ty element insertions, subtelomeric variations and the circularization of one homolog of Chr1 (Heasley and Argueso, 2022). The phasing of polyploid genomes has recently received attention (He *et al*., 2018), with two new long-read methods, WhatsHap polyphase (Schrinner *et al*., 2020) and nPhase (Abou Saada *et al*., 2021) that complement existing phasing methods tailored for diploids containing only two haplotypes (Edge *et al*., 2017; Garg *et al*., 2018; Koren *et al*., 2018; Luo *et al*., 2021; Patterson *et al*., 2015).

This progress illustrates the importance of the technological advances made in DNA sequencing methods and the remarkable developments in computational tools. Among the latter, variation graphs would provide the natural data structure to represent, combine, browse, and query the genomic variation across a population of individuals, thus overcoming the reference bias (Eizenga *et al*., 2020). Another advantage brought by contiguous assemblies is that SV detection across multiple strains can now be achieved by pairwise reciprocal Whole Genome Alignments (WGA) between target genome assemblies and the reference genome (Goel *et al*., 2019; Nattestad and Schatz, 2016; O’Donnell and Fischer, 2020). In *S. cerevisiae*, the most recent long-read datasets containing 8 strains employed a WGA-based SV detection tool, MUM&Co (O’Donnell and Fischer, 2020), and discovered on average 330 SVs per genome (Bendixsen *et al*., 2021). Compared to previous SV detection efforts, this is three times more than the average across other publications with long-read datasets and SV detection based on more classical mapping methods.

Population genomics is entering this new era where numerous contiguous genome assemblies are available, the detection of SVs is more efficient and more complex genomes and their haplotypes are also accessible. Under these conditions, it seems that the concept of reference genome has, or is soon to have, outlived its purpose and will inexorably be replaced by a new paradigm in the form of a pan-genome Reference Assembly Panel (RAP), consisting in the combination of multiple contiguous reference-like assemblies, that would faithfully represent the genetic diversity of the species. Here, we generated the first rigorous RAP using the species *S. cerevisiae* (ScRAP) comprising 142 contiguous genome assemblies that deeply sample the genomic space of the species both in terms of genetic divergence, ploidy, heterozygosity as well as geographic and ecological origins of the strains. In addition to generic haploid assemblies, the ScRAP contains both complete diploid and polyploid phased assemblies. This unprecedented resource allowed us to uncover the structural diversity of genomes across the entire population. We have detected a total of more than 36,000 SVs resulting from over 4,800 different large-scale mutational events. The SV catalog comprises CNVs such as insertions, deletions and duplications but also balanced rearrangements including inversions and translocations. Additionally, The ScRAP revealed the evolutionary dynamics of complex regions that classically remain unassembled in short read-based population genomics projects. These include multicopy tRNA gene families, repeated regions containing Ty elements, subtelomeric regions including the X and Y’ elements, horizontally acquired regions as well as the repetitive telomere sequences located at the termini of the chromosomes, thereby exemplifying the benefit of transitioning from the use of a single reference genome to the use of a large panel of telomere-to-telomere genome assemblies.

## RESULTS

### The ScRAP is a large panel of telomere-to-telomere genome assemblies in *S. cerevisiae*

The ScRAP includes 142 strains that cover the broad geographical and ecological distributions of the species as well as various ploidy and heterozygosity levels (Fig. 1A and B, Supp. Table 1). The full panel is composed of 3 distinct datasets. (i) 100 newly sequenced and *de novo* assembled genomes. These 100 strains were sampled in all previously defined phylogenetic clades (Peter *et al*., 2018) and comprised genomics features such as large SVs, horizontal gene transfers (HGTs) and introgressions (Supp. Table 2). (ii) 18 re-assembled genomes using previously available raw Nanopore read data (Istace *et al*., 2017). (iii) 24 publically available complete genome assemblies, including the S288C reference genome (Bendixsen *et al*., 2021; Berlin *et al*., 2015; Czaja *et al*., 2020; Goffeau *et al*., 1996; Jenjaroenpun *et al*., 2018; Shao *et al*., 2018; Yue *et al*., 2017; Zhang and Emerson, 2019), Supp. Fig. 1). The ScRAP consists of the 142 haploid or collapsed assemblies (one per strain) as well as 55 haplotype-resolved assemblies comprising two phased assemblies per heterozygous diploid (21 strains) and one haplo-phased assembly per heterozygous polyploid (13 strains), totaling 197 nuclear genome assemblies (Table 1, Supp. Table 1). The ScRAP also contains 135 mitochondrial chromosome assemblies consisting of 114 *de novo* assemblies and 21 publicly available mitochondrial assemblies (Table 1, Supp. Table 3).

**Figure 1:**
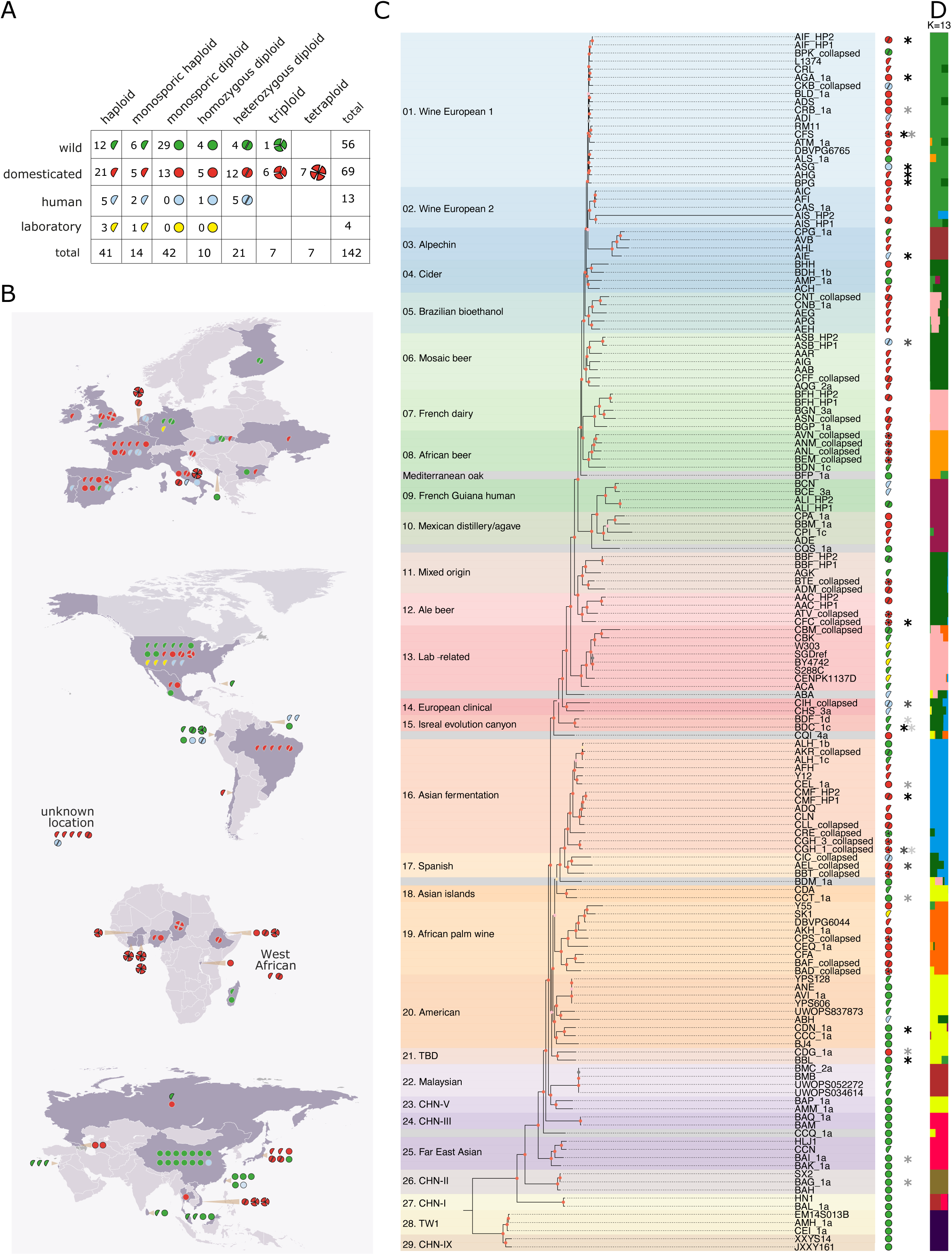
The *Saccharomyces cerevisiae* Reference Assembly Panel. (A) Ecological origin, ploidy and zygosity description of the 142 SCRAP strains. green, red, blue and yellow indicate the ecological origins. Ploidy levels and zygosity are symbolized by the shapes of the symbols. The haploid category contains both natural and engineered (ΔHO) strains. All triploid and tetraploid strains are heterozygous with the exception of the homozygous triploid strain isolated in the USA. (B) Geographical origin of the isolates. Colors and symbols are identical to A. (C) Phylogenetic tree based on the concatenated protein sequence alignment of 1,618 1:1 orthologs. The tree was rooted by including 23 other *Saccharomyces* species used as outgroup (not presented in the figure). The symbols on the right recall the ecological origin, ploidy and zygosity of all isolates. The presence of aneuploid chromosomes is labeled by an asterisk with varying gray levels discriminating between several cases relatively to the 1,011 genome survey (Peter *et al*., 2018): black, previously detected; dark gray, not previously detected; middle gray, previously absent and newly gained; light gray, previously present but newly lost. (D) Genetic ancestry of the population as defined by running ADMIXTURE with k=13.

The number of contigs/scaffolds per nuclear genome assembly reveals high contiguity for all haploid/collapsed assemblies (Supp. Fig. 2A, Supp. Table 1). Additionally, 14 phased diploid assemblies of our newly sequenced genomes show an unprecedented level of contiguity with an average of 25 scaffolds per haplotype. Seven of them are completely assembled into a single telomere-to-telomere haplotype-phased scaffold per chromosome. Based on genome size, the newly sequenced genomes are the most complete, being on average 120 kb and 80 kb larger than the re-assembled and publically available genomes, respectively (Supp. Fig. 2B). The number of chromosomes fully assembled from telomere to telomere is also higher in the newly sequenced genomes as compared to the two other datasets (Supp. Fig. 2C). Noticeably, 47/100 haploid/collapsed assemblies include terminal telomeric repeats at all 32 chromosome ends while only 3/24 public assemblies (12.5%) and none of the 18 re-assembled genomes have all 32 telomeres. The SGD reference genome (S288C) only contains 21 telomeres despite the dedicated effort to reconstruct them (Louis and Borts, 1995). Phased assemblies also approach a telomere-to-telomere completion level with a maximum of 60/64, 76/96 and 101/128 telomeres respectively for the diploid, triploid and tetraploid assemblies. The estimation of the mean telomere length per genome also confirms that the 100 *newly sequenced genomes* are the most complete of all with an average size of 340 bp as compared to 215 bp and 93 bp for public and re-assembled genomes, respectively (Supp. Fig. 2D). Finally, the genome-wide sequence identity with the SGD reference genome is similar across all assemblies, ranging between 98% and 100% with an average of 99.2% (Supp. Fig. 3).

### The ScRAP provide reference-quality genomes across the entire phylogeny

We built a phylogenetic tree based on the concatenated protein sequence alignment of 1,618 1:1 orthologs conserved across all 142 strains of the ScRAP as well as in 23 other *Saccharomyces* species used as outgroup species to root the tree (Fig. 1C). Additionally, a total of 1,581,350 high quality reference-based SNPs were detected across the panel and used to build a second phylogenetic tree. The two tree topologies are similar (Clustering Information Distance of 0.18), differing mainly by local alternative branching of terminal nodes, except for one clinical isolate YJM454 (ABH) that switches between two different positions (Supp. Fig. 4). We analyzed the genetic ancestry of the population by running ADMIXTURE (Alexander *et al*., 2009) on the SNP dataset. The genetic composition of the YJM454 (ABH) strain shows a mixed origin between the ‘American’ and the ‘Mosaic beer/cider’ clades that explains its phylogenetic positioning instability (Fig. 1D). The general topology of both trees is also consistent with previous phylogenetic reconstructions. The ScRAP covers all main phylogenetic clades as they have been previously defined from large population genomics studies (Duan *et al*., 2018; Peter *et al*., 2018), thereby providing reference-quality genomes across the entire species phylogenetic diversity.

Noticeably, the ‘Alpechin’ lineage branches as a sister taxon to the ‘Wine European 2’ clade within the larger population of wine European strains. However, the Alpechin clade has a unique genomic composition, different from that of the wine European strains as it carries abundant *S. paradoxus* introgressions (D’Angiolo *et al*., 2020; Peter *et al*., 2018). The ortholog-based tree brings notable improvement in clade definition by providing reliable phylogenetic structure to previously loose clusters called mosaic regions 1, 2 and 3 (Peter *et al*., 2018) and to isolated branches (Fig. 1C). We defined three new clades called ‘Cider’, ‘Lab-related’ and ‘Spanish’ that comprise isolates previously belonging to the mosaic regions. The existence of these new clades is further supported by their inferred genetic ancestry (Fig. 1D). We also merged the ‘sake’ and ‘asian fermentation’ clades into a single ‘Asian fermentation’ group as well as the ‘North America oak’ and the ‘Ecuadorian’ clades into a new ‘American’ clade. In these two cases, the groupings are consistent with shared geographical origins and their genetic ancestry (Fig. 1B and D). The American clade contains a Chinese strain (BJ4) from the CHNVI/VII clade originally defined in (Duan *et al*., 2018). The genetic relatedness between the American and China VI/VII strains support a recent shared history that likely postdates the main out-of-China event that founded the rest of the world population (Peter *et al*., 2018). The two alternative scenarios of China VI/VII that recently migrated to the Americas or American strains that have re-entered China remain to be defined. One clade containing only two strains isolated from a grape berry in far east Russia (CDG) and from a beetle in Bulgaria (BBL) remains undefined but the strains present an ADMIXTURE genomic composition comparable to that of the American strains (Fig. 1D). The root of the ortholog-based tree groups together the two most diverged Taiwanese I and the Chinese IX wild populations, which is consistent with the East Asian geographical origin of the species (Duan *et al*., 2018; Lee *et al*., 2022; Peter *et al*., 2018; Wang *et al*., 2012).

We leveraged our telomere-to-telomere haplotype-resolved diploid assemblies to investigate the evolutionary histories of the two haplotypes. For all phased genomes, the two sister haplotypes remain always grouped together in the phylogenetic tree and share the same admixture population structure (Fig. 1C). We checked that this was not an artifact resulting from the random assortment of homologous chromosomes within each phased assembly by reconstructing single-chromosome phylogenies and ADMIXTURE structures. All pairs of chromosome-specific haplotypes from the same isolate always branch together showing that both haplotypes in a genome do not result from a recent admixture between different lineages but share a common phylogenetic origin. The most striking difference that we found between two haplotypes from the same isolate concerns the wine European MC9 strain for which the branch length of the haplotype 2 (AIS_HP2) is disproportionately longer as compared to all other terminal branches (Fig. 1C). This was due to chromosome-scale introgressions in this genome where the two homologues of the chromosomes VI and VII originate from two highly diverged species (see below).

We applied a strict molecular clock model to the ScRAP dataset to time the major founding events of the species history (see STAR METHODS). In brief, we used 100 and 365 generations per year as lower and upper bounds (D’Angiolo *et al*., 2020) and mutation rate 2.31 x 10^-10^/bp/generation as estimated from homozygous and heterozygous diploid mutation accumulation lines (Tattini *et al*., 2019). Consistent with previous estimates, *S. cerevisiae* has splitted from its sister species *S. paradoxus* 5.7-1.7 MYA (Supp. Table 4). We estimated that the first split of the most diverged lineage (CHN-IX - TW1) from the species last common ancestor occurred 680-180 KYA. The species’ origin was followed by a single out-of-China event that founded the rest of the world population 290-80 KYA. Multiple independent domestication events occurred likely from wild populations that were established in the north hemisphere. The Wine/European lineage separated 55-15 KYA from the wild Mediterranean oak population, which likely represent his wild ancestor (Almeida *et al*., 2015). All the Wine/European strains shared a common ancestor 43-11 KYA and are nowadays characteristic of winemaking activities across the world.

### The SV catalog covers a large fraction of the entire structural diversity

We identified a total of 36,459 SV calls by pairwise whole genome alignments against the S288C reference genome (STAR METHODS). These calls originated from 4,809 independent large-scale rearrangements that are shared across the other 141 strains.(Table 1, Supp. Table 5). This catalog comprises CNVs such as deletions, insertions, duplications and contractions of repeated sequences (> 50 bp), as well as copy-neutral rearrangements including inversions (> 1 kb) and translocations (> 10 kb, Fig. 2A). We estimated the total SV diversity that is expected at the species level by randomly re-sampling the pool of strains several times and then plotting the average number of non-redundant SV found on each sample. We predict that the species-wide SV inventory would contain approximately 6,000 non-redundant SVs (Fig. 2B, Table 1). Our present SV catalog therefore covers a large fraction (80%) of the estimated species structural diversity.

**Figure 2:**
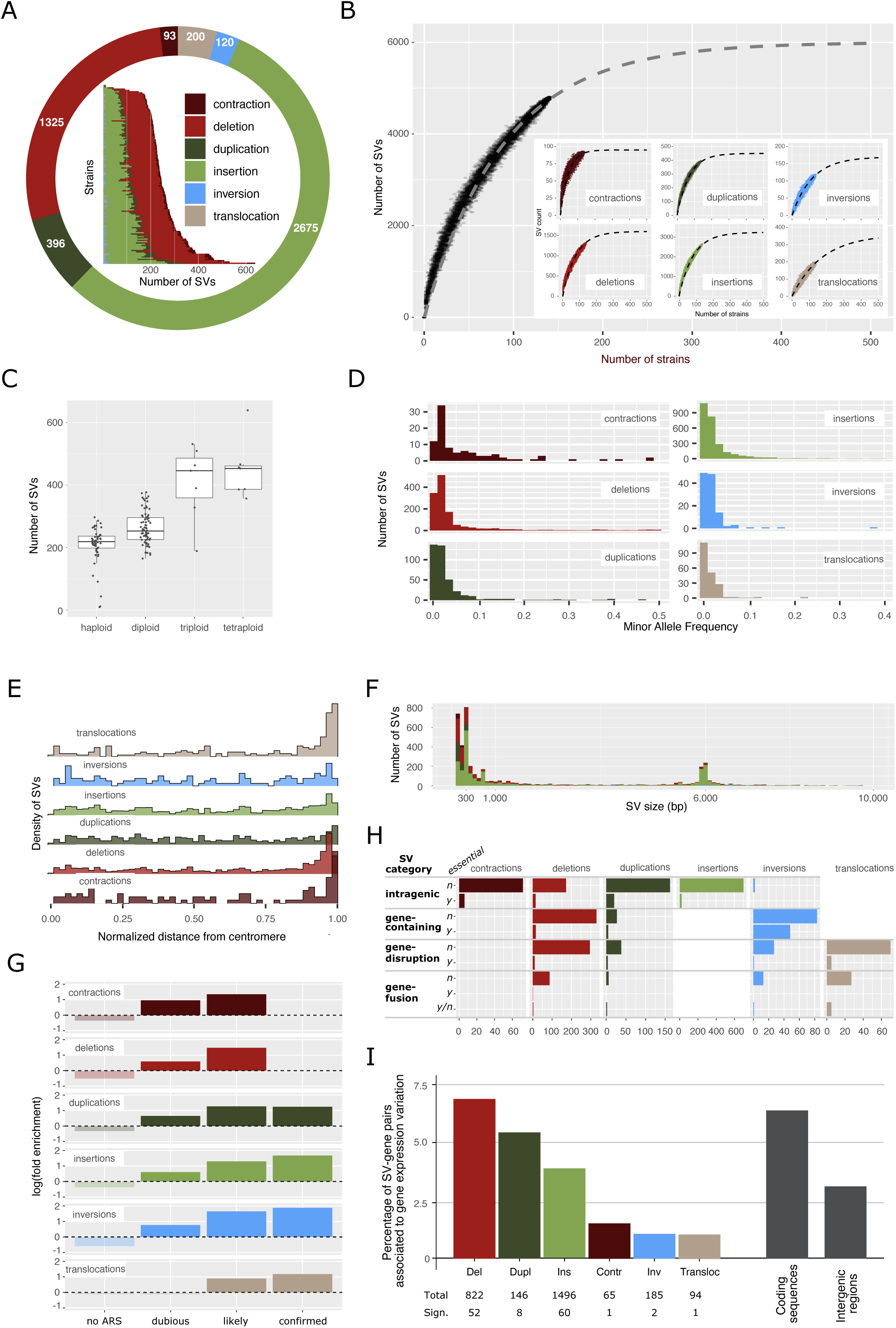
(A) Repartition of the non-redundant set of 4,809 SVs. The outer donut plot indicates the number of SV of each type. The inner bar plot shows the repartition of SV among the 142 strains. (B) Rarefaction curves showing the evolution of the number of non-redundant SV as a function of the number of sequenced strains. A set of 1-141 strains (increasing by increments of 2) was randomly selected 100 times and the number of variants were counted for each set. This was done for all SVs and separately per SV type (inset plots). (C)Number of validated SV per strain split by ploidy. (D) The allele frequency shows how SVs are shared among the strains. (E) Relative density of SV positions along all chromosomal arms split by SV type. The values of 0 and 1 represent the relative positions of the centromeres and telomeres, respectively. (F) Distribution of SV sizes split by type. The colors attributed to the different SV types are as in the other panels. (G) Association between SV and ARS/ORIs. The fold enrichments correspond to the ratio between the proportion of breakpoints associated with a given ORI type and the proportion of the genome covered by the same ORI type. (H) Impact of each SV type on the evolution of the gene repertoire. “Intragenic” means that SVs are fully included within genes. “Gene-containing” means SVs that contain at least one entire gene. “Gene disruption” corresponds to SVs having one breakpoint located within a gene and the other breakpoint in an intergenic region. “Gene fusion” indicates cases where the two SV breakpoints lie with coding sequences. In the *essential* column, *n* (*no*) and *y* (*yes*) mean not essential and essential genes, respectively. (I) Percentage of SV-gene pairs that showed a significant expression change in the presence and absence of the SV depending on the SV type and the location of the SVs, *i.e.* the coding or non coding regions. The numbers at the bottom indicate for each SV type the total number of SV-gene pairs and the number of pairs showing a significant expression difference in the presence or absence a given SV (Ins for insertions, Del for deletions, Inv for inversions, Dup for duplications, Transloc for translocations, and Contr for contractions).

Importantly, phasing heterozygous genomes added a large number of SVs that would have remained undetected using only collapsed assemblies. In the absence of phased assemblies, collapsed genomes would have detected only 67% of all calls for any phased strain, compared to 88% from phased assemblies in the absence of collapsed. Alternatively stated, on average 33% of calls for phased strains were only validated due to phased assemblies (Supp. Fig. 5A). Because SV validation requires a call to be found within at least two genome assemblies, the additional calls validated due to phasing correspond more to heterozygous variants due to the disentangling of their otherwise collapsed status (see SV detection in STAR METHODS). Thus, on average 53% of calls detected in phased strains are heterozygous (Table 1, Supp. Fig. 5B), of which 94% and 45% would have been validated using only phased and collapsed assemblies respectively. Notably, both the proportion of calls validated only in the presence of phased genomes and the proportion of heterozygous variants increases with ploidy.

There is a median of 240 SVs per strain with a range of 10-639 (Table 1, Supp. Table 6). The two strains with the lowest number of SVs correspond to another assembly of the S288C reference strain and its close derivative the BY4742 strain, with 10 and 12 SVs, respectively. The most rearranged genome comprising 639 events is found in YS8(E) (BTE), a mixed origin, highly heterozygous, tetraploid. The second highest number of events is 531 in FTPW4 (CPS), an african palm wine, highly heterozygous, triploid (Fig. 2A). The median number of SV increases with ploidy, from 219 SVs in haploids to 453 in tetraploids (Fig. 2C). The number of SVs does not differ between domesticated and wild isolates (Wilcoxon, P-val=0.53). Deletions and insertions are the most frequent types of SVs with about 100 events per strain, followed by duplications and contractions globally represented 10 to 20 times per strain, translocations and inversions are rarer with only a few occurrences on average per strain (Supp. Table 6). MAF shows that contractions are more commonly shared than all other SV types. However, most SVs are present at low frequencies in the population, with 34% of the events being found in a single genome and 91% with a minor allele frequency (MAF)□<□0.1 (Fig. 2D). This suggests that SVs would be mostly deleterious and that negative selection would prevent most of them from spreading in the population.

### SVs are enriched in specific chromosomal locations, genetic elements and breakpoint-associated sequences

We looked at the repartition of SVs along the chromosomes by plotting the distance of SV breakpoint to chromosome ends and centromeres. All types of SVs, but the inversions, are primarily confined to subtelomeric regions (Fig. 2E), in accordance with the high evolutionary plasticity of these regions (Yue *et al*., 2017). Insertions more frequently contain repetitive sequences (82%) compared to deletions, duplications and inversions (41-47%) (Supp. Fig. 6). The distribution of event sizes, excluding translocations, shows that small SVs are the most frequent with 58% and 9% of the events being smaller than 1 kb and larger than 10 kb, respectively (Fig. 2F). This distribution shows two clear peaks around 300 bp and 6kb for deletions, insertions and inversions corresponding to solo-LTRs and full-length Ty elements. The mobility of Ty elements directly accounts for 59% of all insertions (1,571 events) but only 16% of deletions through inter-LTR recombination (218 events). This difference is understandable, as there is only a limited number of Ty elements that are present in the reference genome that can be interpreted as a deletion when absent from other genomes. Interestingly 19% and 8% of the duplications (n=74) and contractions (n=7), respectively, also result from the direct movement of Ty elements that occurred in tandem. Altogether 39% of all SVs result from the insertion and deletion of Ty elements.

We examined breakpoint sequences found at the junction of the different SV categories. In order to remove reference-bias, we analyzed each event separately in each strain based on its *de novo* assembly coordinates and the overlapping genetic elements listed in the strain annotation files. Following this we ran an enrichment test for each genetic element. We found a clear enrichment of repeated sequences (LTRs, Tys, tRNAs, Y’ and X elements) at the junction of all types of SVs as well as a significant underrepresentation of CDS within breakpoints (Supp. Fig. 7). Interestingly, we found a significant association between replication origins (Autonomously Replicating Sequences, ARSs) and SV breakpoints. Furthermore, we extracted all ARSs from ORIdb (Siow *et al*., 2012), including dubious and likely sites and showed that the ARS-SV association is greater as the likelihood of the ORI being used increases (Fig. 2G).

### SVs play a major role in the evolution of the gene repertoire

It is clear that SVs have an important evolutionary potential given their intrinsic capacity to modify the gene repertoire. We found that nearly 40% of the SVs (1876/4809) directly impacted protein coding gene sequences (Table 1), and these SVs are not those involved in the insertion and deletion of Ty elements. Interestingly, this proportion drops to 3% when considering only the essential genes (160/4809), as defined by their associated ‘inviable’ phenotype in SGD (1,117 essential genes in total). We defined 4 different categories depending on the relative arrangement of the SV breakpoints and the impacted coding sequences (Fig. 2H). The most frequent case is by far the situation where both breakpoints of a given SV lie within the same gene sequence (this by definition excludes translocations). We found 1170 such cases of intra-genic SVs, majoritarily corresponding to insertions and to a lesser extent to deletions and duplications. Most contractions of repeated sequences also belong to this category as 78 of them, out of a total of 93 events, fall within coding sequences. It is difficult to make predictions on the functional outcome of intra-genic SVs as each event can disrupt, or not, its corresponding coding sequence depending on its size and phasing relative to the original reading frame. Anyhow, this SV category has a strong potential to increase the diversity of the gene repertoire. The second category gathers all SVs that contain at least one entire gene (excludes insertions and translocations, Fig. 2H). We found 508 cases of such gene-containing SVs, corresponding to 345 deletions, 84 inversions and 27 duplications containing on average 5, 30 and 2 genes, respectively. In total, the 345 deletions comprised 525 different genes that have been fully deleted in at least one haplotype. The manual review of the 29 deleted genes that are described as essential revealed that they are in fact nonessential or are conditionally essential or are found deleted only at the heterozygous state. The two last categories, gene disruption and gene fusion, comprise all SVs for which either one or both breakpoints lie within a protein coding gene (excludes insertions). Note that these two categories are not mutually exclusive with the previous one as a given event can both contain entire genes and disrupt or fuse other genes at its breakpoints. We identified 450 cases of gene disruptions that produce gene truncations by merging the internal part of a gene with an intergenic region. We also found 145 cases of gene fusion where both borders of a given SV interact with different genes. These events have the capacity to create new chimerical genes. Note that fusions happened between essential and non-essential genes in 11 cases. Deletions are the most prevalent SV type in these categories. Surprisingly, about half of the translocations (98/200) resulted in gene disruption (n=71) or fusion (n=27) at their breakpoints, while it is generally assumed that translocations occur mostly between dispersed copies of transposable elements.

Altogether, we identified 1,698 complete gene deletions and duplications, as well as 1,513 gene structure alterations at the origin of new gene sequences that can significantly expand the gene repertoire of the species.

### SVs impact gene expression nearby their breakpoints

The assumption that SVs play a major role in phenotypic variation is now well recognized and a large number of studies have provided clear specific examples but there is no comprehensive description of the impact of SVs at a population-scale. In this context, SVs can impact gene expression and more precisely the expression of genes located nearby. SVs can potentially alter or influence gene expression by impacting the sequence of the ORFs (Open Reading Frame), modifying their copy numbers as well as changing the *cis*-regulatory elements. By leveraging a recent survey that generated the transcriptome of more than 1,000 *Saccharomyces cerevisiae* isolates (Caudal et al. submitted), we explored the relationship between gene expression and the exhaustive catalog of SVs we defined here. For 51 out of 142 isolates, we obtained the transcriptome, consisting of the expression levels for 6,445 transcripts and a total of 1,876 SVs, encompassing a similar proportion of the different SV types than the entire dataset (Supp. Fig. 8A).

To explore the effect of SVs on gene expression, we first defined the entire set of SV-gene pairs, representing all the SVs located in the coding sequence or in the intergenic region neighboring a gene (see STAR METHODS). Overall, we captured 2,808 SV-gene pairs with more than half of the pairs involving insertion and deletion events. We then compared the expression of genes associated or not with a given SV in our population (Supp. Fig. 8B, see STAR METHODS). We found that 124 SV-gene pairs (4.4%) (Supp. Table 7), encompassing a set of 97 unique SVs, showed significant differential expression changes (Table 1). This impact seems to be subtle but it is important to mention that the transcriptomic data were obtained for cultures in complete medium. Also, here we are only looking at direct *cis* effects. Interestingly, this proportion is variable depending on the type of SV considered (Fig. 2I). While more than 5% of pairs involving deletions and duplications showed significant expression changes, this was only the case for about 1% of pairs involving inversions, translocations and contractions. These results demonstrate a variable impact on gene expression depending on the type of SVs, with deletions, duplications and insertions having a greater impact.

We then explored the difference in impact between SVs located in coding sequences and those that could affect regulatory regions. Due to the structure as well as the prevalence, only the insertions and deletions were considered in this analysis. In total, 7.3% (60 out of 815) affecting coding sequences are associated with significant differences in expression, mainly by reducing or suppressing expression (Fig. 2I). By contrast, only 3.1% (23 out of 726) pairs associated with SVs present in non-coding regions were detected as significantly impacting gene expression.

Overall, we found over one hundred SV-gene pairs that exhibit significant impact on gene expression and could broadly influence quantitative trait variation. Interestingly, we also highlighted that specific SVs (i.e., insertions, deletions and duplications) as well as SVs affecting the coding sequences have a stronger impact on gene expression.

### SVs and SNVs follow distinct evolutionary trajectories

We built a phylogenetic tree using the 36,459 SVs identified among the 142 strains and compared its topology to that of the SNV-based tree (Fig. 3A). The two trees show a globally conserved organization (Clustering Information Distance of 0.44) with the most diverged isolates corresponding to the Chinese and Taiwanese strains and most clades, as defined from the ortholog tree (Fig. 1C), remaining individualized and recognizable. However, the relative position of the clades is poorly conserved between the two trees. For instance the two sisters clades, 10. Mexican distillery/agave and 09. French Guiana human, remain grouped together in both trees but their position switched from a more external node in the SNVs/INDELs tree to a more internal node in the SV tree, close to the 18. African palm wine clade. These discrepancies could be due to a less reliable phylogenetic reconstruction in the SV-based tree as indicated by lower confidence scores, only 56% of the nodes are supported by confidence values higher than 0.95 as compared to 93% in the SNV-based tree (Supp. Fig. 9). This is likely due to a much lower number of distinct events (4,809 SVs *vs* 1,581,350 SNPs) as well as low allele frequencies for SVs (91% with MAF<0.1). However, in some cases, the evolutionary signal brought by SVs is meaningful. For instance, the BJ4 strain, which was isolated from China and previously located in a Chinese clade (Peter *et al*., 2018), now branches in a more internal region of the tree along other Chinese strains (Fig. 3A) while it clusters within, or next to the American clade containing primarily north American oak samples in both the ortholog-based and SNV-based trees (Fig. 1C, Fig. 3A). Estimation of individual genetic ancestries by ADMIXTURE (Alexander and Lange, 2011) using SVs as markers also confirm the phylogenetic clustering defined by the SV-tree. For example, the DBVPG1841 (BPG) strain shares the same ancestry as the CBS7964 (AEH) strain, supporting their clustering in the SV-based tree, while these two isolates are located in different clades in both the ortholog and SNP-based trees. Several other examples are visible on Fig. 3A. In conclusion, it appears that the two types of polymorphisms SNVs/INDELs and SVs give different signals and that the signal brought by SVs is less precise but complementary to that of SNVs/INDELs for evolutionary reconstruction.

**Figure 3:**
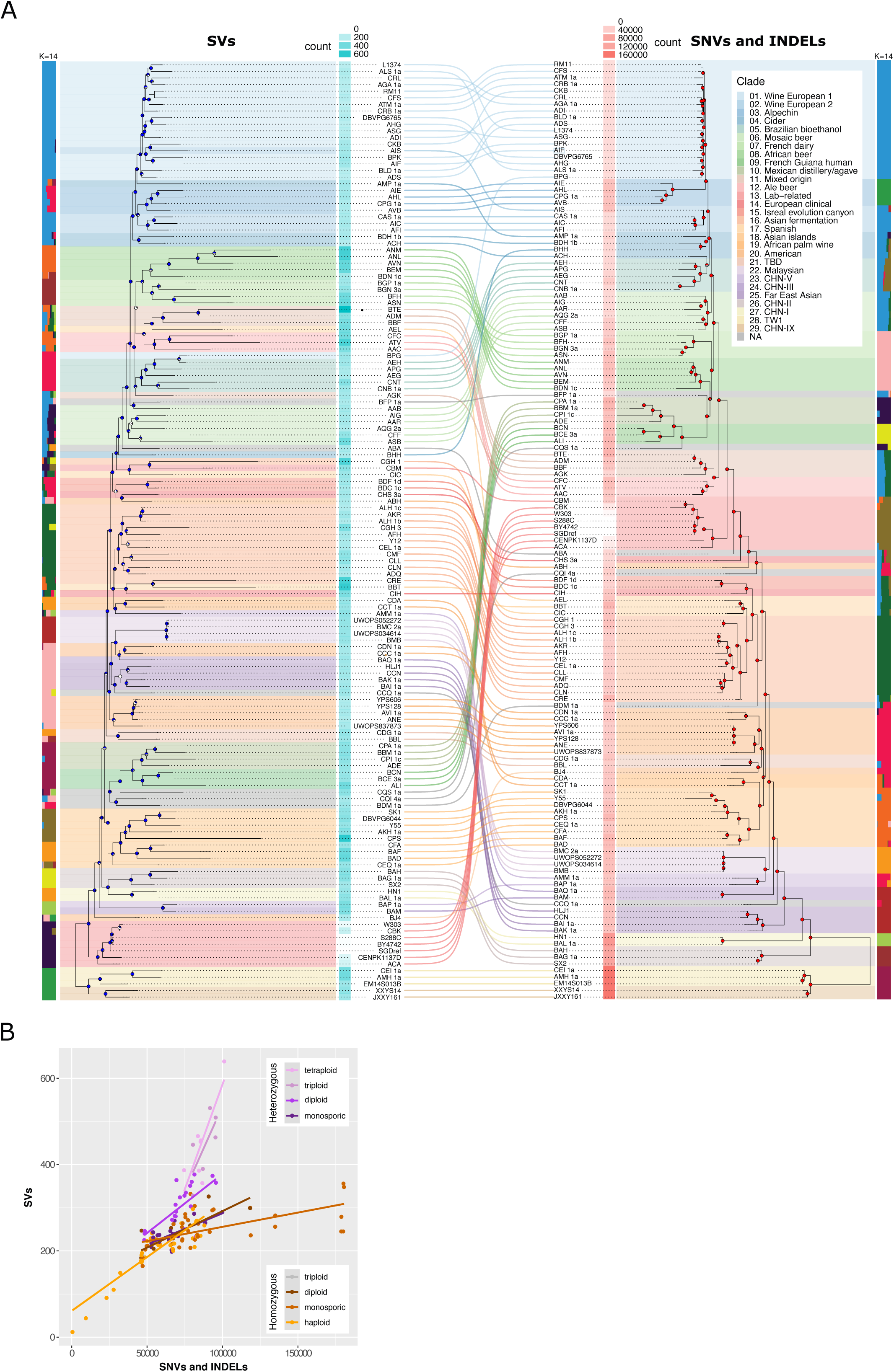
A) Comparison between the SV-based (left) and SNV/INDEL-based (right) phylogenetic trees. The genetic ancestry of each strain was inferred by ADMIXTURE using SVs (k=16) and SNVs/INDELs (k=14) and the number of SVs and SNVs/INDELs per strain is presented as cyan and coral heatmaps, respectively. (B) Accumulation of SVs as a function of SNVs and INDELS. The definition of heterozygous and homozygous strains is solely based on the SNVs/INDELs data. The categories ‘Heterozygous monosporic’ and ‘Homozygous monosporic’ correspond to monsporic isolates derived from the sporulation of heterozygous and homozygous parental diploid strains, respectively.

The relative branch lengths are also different between the two trees (Fig. 3A), suggesting independent accumulation of the two types of mutational events. We plotted the number of SVs as a function of the number of SNVs/INDELs for each strain and found a clear positive correlation between the two variables. However, heterozygous and homozygous genomes show different patterns (Fig. 3B). For a given number of SNVs/INDELs, the number of SVs is systematically higher in heterozygous than in homozygous genomes. As the distinction between homozygous and heterozygous genomes was solely based on SNVs, we reasoned that the lower number of SVs in SNV-based heterozygous genomes could possibly be due to a lack of detection of heterozygous SVs because SNV-based homozygous genomes were not phased and therefore we could expect only half of the heterozygous SVs to be detected. In order to test this hypothesis we ran Sniffles (Sedlazeck *et al*., 2018), a read-based SV detection tool, which is therefore not prone to missing heterozygous variants as compared to assembly based methods with unphased genomes. We found the same general pattern using this method, more SVs relative to SNVs in heterozygous strains, and found no evidence that any significant number of heterozygous SVs were missed in our homozygous strains (Supp. Fig. 10A). Furthermore, the same association was found using paftools (Li, 2018), another assembly-based SV detection tool using minimap2 mapping as opposed to nucmer (Supp. Fig. 10B). These analyses show that the higher number of SVs found in heterozygotes as compared to homozygotes is not due to a methodological bias and suggest that SVs preferentially accumulate or are better tolerated in heterozygous genomes.

Another striking feature is the association between ploidy and SVs. We previously showed that the median number of SV increases with ploidy (Fig. 2C). It appears here that for a given number of SNVs/INDELs, the higher the ploidy the more SVs (Fig. 3B). This cannot be simply explained by the increasing genome size with ploidy as this is expected to similarly affect both SVs and SNVs/INDELs.

### Complex aneuploid chromosomes comprise large SVs

We identified 26 simple chromosome aneuploidies affecting 18 out of the 142 isolates of the panel (Supp. Table 8). This represents a lower proportion compared to what was reported in the 1,011 population (13% *vs* 19% of the strains, respectively). This difference is probably due to the selection of preferentially euploid strains when we chose the ScRAP isolates. Interestingly, thanks to the long-read sequencing and *de novo* assemblies, we discovered more complex aneuploidies where aneuploid chromosomes comprise large SVs such as inter-chromosomal translocations or large insertions or deletions. We identified 8 such cases of complex aneuploidies in 7 strains, which represents 24% of all aneuploidies found in the panel (Supp. Table 8) and suggests that this phenomenon could be common. Of these complex aneuploid chromosomes, 4 contain translocations, 1 contains both a translocation and an HGT insertion, 2 contain large deletions (∼100 kb), and the structure of the last complex event remains unresolved (in strain PB12 (CIH), Supp. Table 8). We managed to fully resolve the chromosomal organization in 5/7 strains (Fig. 4A). The detailed chromosomal structure in the strain CBS457 (AIF) could not be fully resolved as in addition to a chrXVI_chrIV aneuploid translocation, another complex chrXI_chrXIV aneuploid translocation was identified but its fine structure remained elusive. We confirmed that all 7 complex aneuploidies were already present when the strains were initially Illumina sequenced (Peter *et al*., 2018) but not identified at that time.

**Figure 4:**
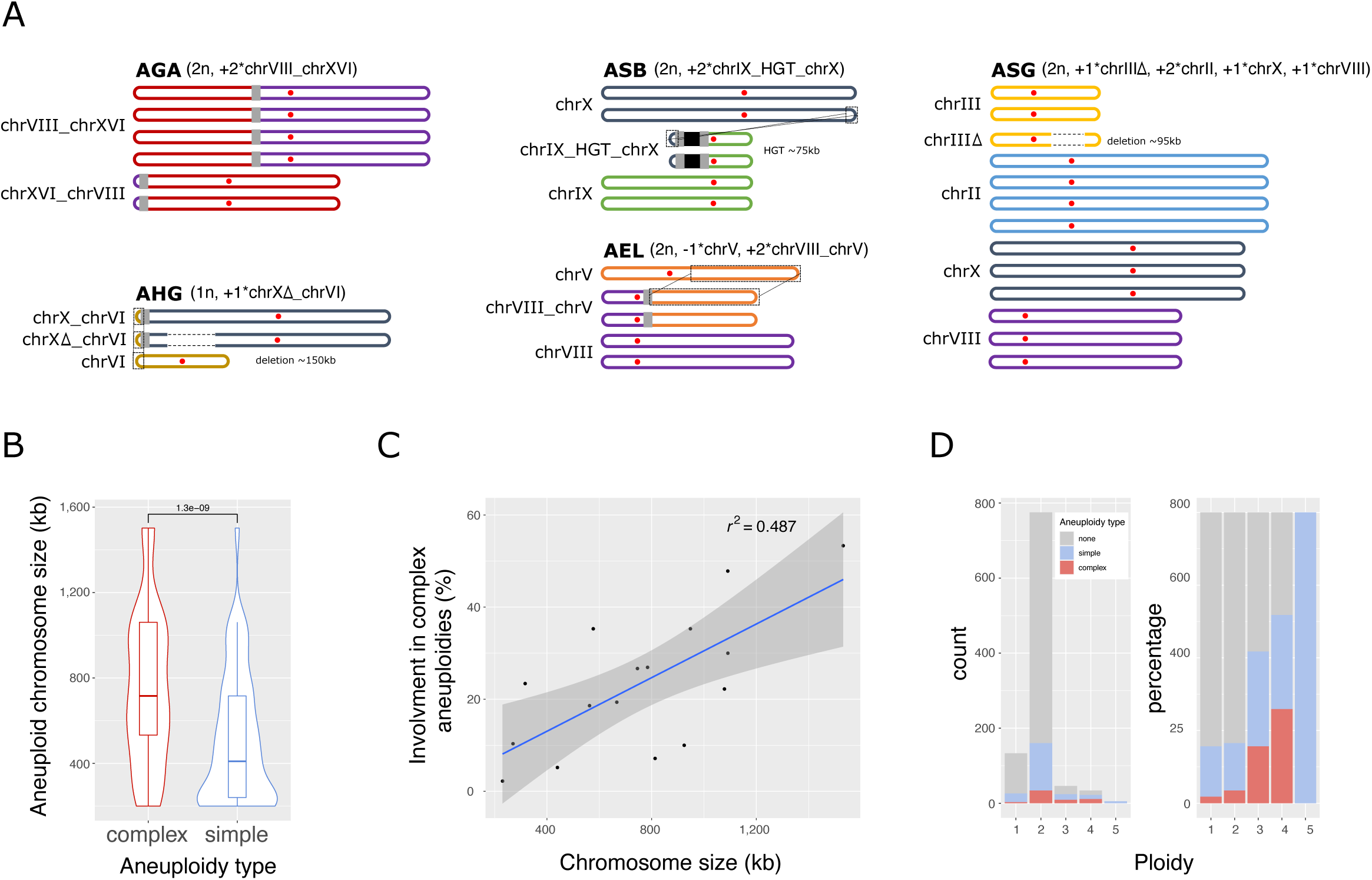
(A) Schematics representing the chromosomal composition of complex aneuploidies. The red dots symbolize centromeres. The grey boxes indicate the translocation breakpoints. Dotted framings show duplicated regions while dotted lines within chromosomes indicate deletions. The black box in ASB symbolizes the HGT region. (B) Distribution of the original sizes of the chromosomes involved in complex *vs* simple aneuploidies. (C) Percentage of chromosome involvement in complex aneuploidies as a function of their size. (D) Number and percentage of euploid, simple and complex aneuploid among 993 analyzed strains.

In order to understand if this complex aneuploidy phenomenon was generalisable to a larger set of strains, the aneuploidy status of 993 strains (84 from (Strope *et al*., 2015) and 909 from (Peter *et al*., 2018)) was re-analysed to detect both simple and complex aneuploidies (see STAR METHODS). The proportion of aneuploid chromosomes estimated to be complex in this larger dataset is between 10-18% (44/423 - 85/464). In terms of strains, 15-23% (36/248 - 59/248) of aneuploid strains are predicted to contain at least one complex aneuploidy (Supp. Table 9, Supp. Table 10). Lower boundaries of 10% of complex aneuploid chromosomes in 15% of the strains are conservative estimates as they only consider Centromere-Related (CR) events *i.e.* regions where the coverage deviation covers a centromere (for instance chrVIII in AGA in Fig. 4A). The upper boundaries of 18% of complex aneuploidies in 23% of aneuploid strains are less conservative estimates as they also include chromosomes with large (>100kb) Non-Centromere Related (NCRs) coverage deviations, if the same strain includes another CR event (for instance chrXVI in AGA in Fig. 4A). It is fair to use the less conservative estimate including NCRs because of the clear association between CRs and both more frequent and large NCRs (Supp. Fig. 11), and also because we characterized several examples of large translocations being at the root of complex aneuploidies (Fig. 4A). False positive NCRs would only correspond to the co-occurrence of both a large segmental duplication (> 100kb) and at least one simple aneuploid chromosome in the same genome and therefore should remain limited. Therefore, if a strain contains an aneuploid chromosome and a large NCR, the NCR chromosome can be considered also as an aneuploid-related chromosome. In summary, this analysis suggests that a large proportion of aneuploid chromosomes (up to 18%) are associated with large SVs at the population scale.

We also found that aneuploidies are highly unstable given that several cases are different between the initial state described in the population survey of 1,011 isolates (Peter *et al*., 2018) and the chromosomal content that we resequenced in this study, with both new gains and losses of aneuploid chromosomes. We found 6 aneuploidies, in 4 strains, that were lost since the initial Illumina sequencing, either by the gain (5) or loss (1) of a chromosome. Two out of the 6 lost aneuploid chromosomes appear to be complex aneuploidies that were previously undetected. Conversely, we also found 7 cases where chromosomes became newly aneuploid since the initial survey, of which 6 occurred in the monosporic isolates. We also noted that 5 cases were not initially reported (Peter *et al*., 2018), although present and 6 have changed their aneuploidy status (5 gained an extra copy and 1 changed from +1*chr1 to -1*chr1). Notably 3/5 aneuploid chromosomes not reported previously, were complex aneuploidies. More globally, the 1,011 survey reported 343 aneuploid chromosomes in 200 strains (Peter *et al*., 2018). Reanalysing the same dataset here, we found 120 additional cases where 35 of them (29%) are made of complex calls, showing that complex aneuploidies were prevalently lacking in initial analysis.

Interestingly, we found that complex aneuploidies involve significantly larger chromosomes as compared to simple aneuploidies (Fig. 4B). Additionally, there is a clear positive correlation between the involvement of the different chromosomes in complex aneuploidies and their size, showing that larger chromosomes are more often involved in complex aneuploidies than the smaller ones (Fig. 4C). Finally, a general pattern of increasing proportion of simple aneuploidies with increasing global ploidy of the strains has been reported (Gilchrist and Stelkens, 2019). In line with this, we also found an increasing proportion of complex aneuploidies with increasing ploidies (Fig. 4D).

### Evolutionary dynamics of complex genetic elements

#### Telomere length at TEL03L is greater in the absence of Y’ element

We characterized the telomere length for each chromosomal end in all the 142 strains of the panel and found globally significantly larger telomeres in our 100 *newly sequenced genomes* as compared to both public assemblies and re-assembled genomes (Supp. Fig. 12). Some methodological limitations could have resulted in a systematic telomere length underestimation in previous studies. Therefore, all subsequent analyses on telomere length are restricted to the 100 *de novo* genome assemblies in order to avoid any size underestimation bias. The mean telomere size varies by a factor of 4 across the different isolates, from 166 bp in the CBS2183 wine strain (AFI) to 686 bp in a monosporic isolate derived from the CLIB561 French dairy strain (BGN_3a). Some strains harbor homogeneous telomeres across different chromosome extremities while others harbor a large variance in the trait (Supp. Fig. 12). The average telomere length per strain correlates positively with variance (Supp. Fig. 13A). We found no significant correlation between telomere length and ploidy, heterozygosity or ecology (Supp. Fig. 13B-D).

The determinants of the telomere length diversity at the population level are not fully understood. The model for individual telomere length regulation that is currently accepted in several different organisms is the protein counting model suggesting that the probability of telomere elongation increases when telomere length decreases (Marcand *et al*., 1997, 1999; Teixeira *et al*., 2004). This model implies that all individual telomeres would gradually converge toward the population mean length distribution after a sufficiently large number of divisions and that the overall telomere length distribution would converge to a steady state (Bianchi and Shore, 2008; Xu *et al*., 2013). However, a recent study that used nanopore sequencing to measure individual telomere length reported that in W303, each chromosome end could have a specific and reproducible telomere length distribution that would remain stable over at least 120 generations (Sholes *et al*., 2021). We examined telomere length variation between individual chromosome ends across the population of 100 strains. Despite a globally homogeneous distribution, TEL03L, and to a lesser extent TEL07R, are significantly longer than all other telomeres (Fig. 5A, Supp. Fig. 14). The two same chromosome extremities were also described as the longest in W303, (Sholes *et al*., 2021). The conservation of larger telomere size at TEL03L at the population level suggests that the underlying genetic determinants would be conserved since the species diverged from its last common ancestor at least ∼180 KYA (see above). Therefore, we looked for the presence of cis-acting subtelomeric genetic elements whose presence or absence could possibly contribute to the largest size of TEL03L at the population level. For core X, we found that most ends carry a single copy of the element (67%, 3036/4528) and telomere length is significantly longer at chromosomal extremities devoid of this element (Supp. Fig. 15A and B). This difference however is not visible at TEL03L, showing that the largest telomere size at this end could not be explained by the presence or absence of core X (Supp. Fig. 15C). We found that the TEL03L was specifically enriched in *Ty5* elements, along with TEL11R. Considering all ends together, telomere length was significantly larger when subtelomeres contained a *Ty5* element (Supp. Fig. 15D and E). However, the telomere length specifically at TEL03L is not influenced by the presence of *Ty5*, ruling out the possibility that this element regulates telomere length in *cis* (Supp. Fig. 15F). Finally, we found that TEL03L is depleted in Y’ elements, being actually the poorest of all ends across the population (Supp. Fig. 15G). There are on average 17 Y’ elements per genome. The majority of chromosome ends are devoid of Y’ elements (58% or 2642/4560), while 36% (1660) have a single element (Supp. Fig. 16A). We found two strains that are completely devoid of Y’ elements, one from Ecuador, CLQCA_20-156 (CCC_1a), and one from Nigeria, PW5_b (ADE, Supp. Fig. 16B). Noticeably, the haploid YJM981_b clinical isolate (ADI) harbors an extremely high number of Y’-element (Bergström *et al*., 2014), with 154 subtelomeric assembled copies (full-length + partial) as compared to a median of 16.5 elements in the *de novo* haploid/collapsed assemblies (Supp. Fig. 17, Supp. Table 11). As a result, this strain has the largest genome of all with a genome size that is 920 kb larger than the median genome size (12.89 Mb *vs* 11.97 Mb, Supp. Fig. 18A). When considering all chromosome extremities together, telomere length is not different between Y’ pime-containing and Y’-devoid subtelomeres (Supp. Fig. 15H). However, TEL03L that contain a Y’ have significantly shorter telomeres than the ones devoid of such element (Fig. 5B) and this trend is unique to TEL03L among the 32 chromosome extremities (Supp. Fig. 19). This finding suggests that the effect of the sequence that promotes the formation of longer telomeres at TEL03L would be specifically buffered by the presence of a Y’ element at this extremity.

**Figure 5:**
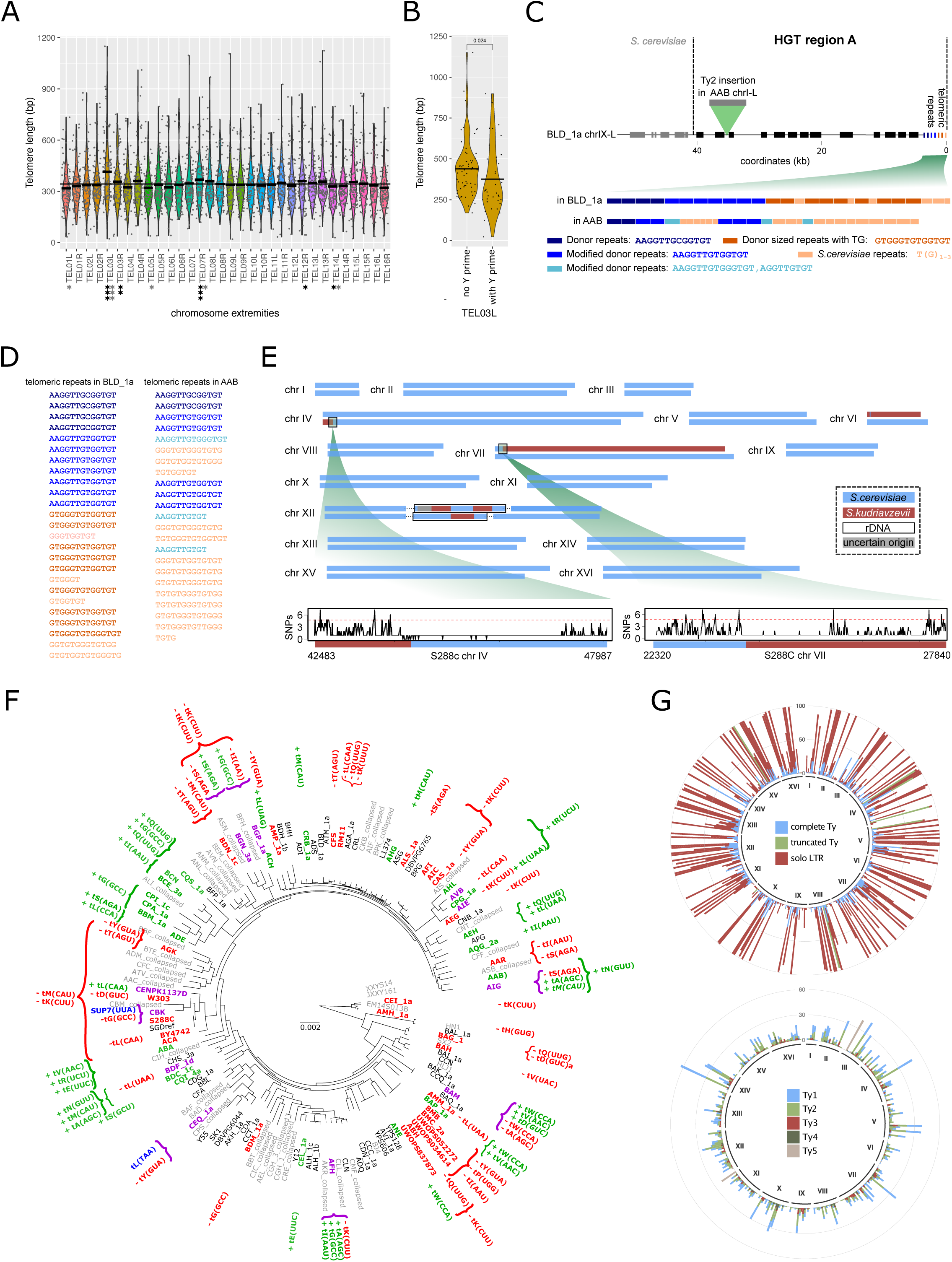
(A) Individual telomere length across the 100 newly sequenced genomes. The crossbars within each violin plot indicate individual mean telomere lengths and the horizontal line shows the global mean telomere length across all ends. Black and gray stars represent the rank product and the rank sum variant statistical tests with 1, 2 and 3 stars corresponding to P-values < 0.05, 0.01 and 0.001, respectively.(B) Distribution of telomere length on TEL03L as a function of Y’ elements. The crossbars within each violin plot indicate individual mean telomere lengths.(C) Region A was transferred from an unknown *Torulaspora* species and is found at the telomeric ends of two ScRAP isolates. One strain (isolate BLD_1a) contains the complete region, while in AAB the region is interrupted by the insertion of a *Ty2* from *S. cerevisiae*. In both cases, the telomeric repeats are completely assembled. Interestingly, the most inner part of the repeats is extremely close to the *Torulaspora* repeats (AAGGTTGCGGTGT here, AAGGTTG**A/T**GGTGT in *Torulaspora*, (Peska *et al*., 2021)), while the distal portion consists of the conventional *Saccharomyces* (TG_1-3_). (D) Between the two types of telomere repeats, mixed repeats are detected, which gradually transition from *Torulaspora-*type to *S. cerevisiae-*type (e.g. AAGGTTG**T**GGTGT -> **GT**GG**G**TGTGGTGT in BDM). The different colors correspond to the repeat types presented in panel C. (E) Karyotype of the two phased haplotypes of the AIS strain. The origin of the haplotypes was inferred based on similarities between each AIS homologue and the genome of S288C (blue) and *S. kudriavzevii* CR85 (dark red) revealing introgression of an almost complete copy of *S. kudriavzevii* chromosomes VI and VII, the terminal part of the left arm of chromosome IV and rDNA sequences on both haplotypes. Both breakpoints on chromosomes VI and VII occur in regions that have low sequence divergence compared to the genome wide average (1 SNPs every 4.78 bp, red dashed line).(F) tRNA gene gains and losses across the phylogenetic tree. The topology of the tree is the same as in Figure 1. tRNA gene gains and losses are represented in green and red, respectively. Anticodon modifications are written in blue. The names of the strains are colored according to the type of events they underwent, in green for gains, red for losses and in purple when different types of events co-occurred. (G) Chromosomal repartition of all types of transposable elements across the 100 de novo assembled genomes (top). For isolates harboring several types of elements in a single region, the complete Ty is preferentially represented, followed by the truncated Ty. Chromosomal repartition of complete Ty elements across the ScRAP (bottom). Only one element is plotted per isolate. For isolates with several families of complete Ty in a given insertion site, the family found in the reference genome was preferentially represented.

#### HGT regions constitute new telomeres

Multiple HGT events have been reported in *S. cerevisiae* but their mechanistic origin and accurate structure have remained elusive (Legras *et al*., 2018; Novo *et al*., 2009; Peter *et al*., 2018). We searched in the ScRAP all large HGT regions known in *S. cerevisiae* (regions A to G, Supp. Table 12) and characterized their structure and evolution. An emerging shared feature is that they are all localized at telomeres, which implies that they must preserve or restore telomeric sequence and function upon their transfer. By resolving the chromosomal structure of HGT regions we revealed multiple ways to (re)generate functional telomeres.

Region A (40 kb) has been transferred from an undefined *Torulaspora* species (Novo *et al*., 2009) and is present at chromosome IX-L in DBVPG1608 (BLD_1a) and chromosome I-L in CBS422a (AAB) (Fig. 5C, Supp. Fig. 20). Additionally, the region A in AAB has a Ty2 insertion, showing how these regions further evolved after the transfer. We next inspect the telomeric repeats in the distal part of region A and observe internally telomeric repeats of the *Torulaspora* donor species gradually shifting to the classic *S. cerevisiae* TG_1-3_ repeats, with some intermediate repeats with a mixed composition (Fig. 5C and D). This structure suggests that the *Torulaspora* repeats have seeded *de novo* telomere addition by telomerase to reconstitute a functional *S. cerevisiae*-like telomere.

Region B originates from *Zygosaccharomyces parabailii* (Galeote *et al*., 2011; Peter *et al*., 2018). We detected region B in 28 strains (Supp. Fig. 20). We assembled a complete copy (98 kb) and closely inspected its boundaries. First, we observed a local region of homology (75%) including several tracts between 10 and 20 bp of complete identity near the insertion site (Supp. Fig. 21A). Such homology is highly unexpected given the extreme sequence divergence of the two species (>30%) and suggests that HGTs could possibly be driven by sequence homology, as for introgressions, their insertion sites might be constrained by local divergence. At the telomere side, we observed the pure *S. cerevisiae* telomeric repeats flanked by the Y’ element. The Y’ element origin is still elusive and so far has only been found in species belonging to *Saccharomyces* complex closely related to *S. cerevisiae* (Liti *et al*., 2005). Surprisingly, we observed that the Y’ element is present in the *Z. parabailii* genome and likely derives from a sister *Saccharomyces* species. The strains containing the region B, show both types of Y’ sequence, with a subset of them having the *S. cerevisiae*-type and others the *Z. parabailii*-type (Supp. Fig. 21A). Taken together, these results support a complex multi-step HGT scenario with the Y’ element from a *Saccharomyces* species that initially invaded the *Z. parabailii* genome and inserted at multiple subtelomeres and was secondarily transferred in *S. cerevisiae* together with the flanking region B. It is possible that the primary transfer of Y’ in *Z. parabailii* played some role in the mobilization, insertion and stabilization of the HGT associated chromosome end.

We previously proposed region G as a new large HGT with the peculiarity of being present in a single wild strain isolated from Ecuador from an unknown source (Peter *et al*., 2018). Indeed the *de novo* assemblies confirm the presence of region G at the chromosome XIII-R subtelomere of the haplotype 2 of the CLQCA_20-060 (ALI) strain (Supp. Fig. 20, Supp. Fig. 21B). Similarity search of the protein coding sequence supports an undetermined *Lachancea* species being the region G donor.

Finally, while all the large HGT regions are present at telomeres, complete assemblies illustrate how these regions can evolve and re-localise in the core chromosome regions. This dynamic is perfectly exemplified by the region F (60 kb), which we detected both in long-subtelomeric and short-internal configurations in two unrelated strains (Supp. Fig. 20, Supp. Fig. 21C). The most parsimony scenario is that the long-subtelomeric is ancestral and the short-internal is the evolved state that shortened during internalization.

Overall, although the molecular mechanisms initiating the HGT remain elusive (e.g. uptake of naked DNA or transient hybrids/cytoduction), the complete assemblies have revealed a role for both sequence homology at the insertion site and the requirement for telomere-repeats type at the extremity that likely constrain the landscape of yeast HGTs. While HGTs are enriched in domesticated strains, they are also present in wild isolates (Supp. Fig. 20), suggesting that they can also occur in natural environments and perhaps get amplified under anthropic conditions.

#### Introgression of entire chromosomes from a remote species

Haplotype phasing of the MC9 (AIS) strain, isolated from Vino Cotto in Italy, revealed a hitherto unseen case of chromosome-scale introgression. One complete homolog of chrVI and a nearly complete homolog of chrVII introgressed from *Saccharomyces kudriavzevii*, exemplifying a unique type of karyotype that does not fit with classical cases of hybrid or introgressed genome structures (Fig. 5E). *S. cerevisiae* (Sc) and *S. kudriavzevii* (Sk) hybrids have been reported in wine making environments (González *et al*., 2006), but introgressions have not been observed. While hybrids with different genome structures from highly diverged *Saccharomyces* parents have been reported, blocks of introgressed materials have only been observed between sister *Saccharomyces* species. The lack of introgression from highly diverged species is likely due to extreme sequence divergence suppressing recombination and genetic incompatibilities. However, we investigated the sequence divergence at the recombination breakpoints of chromosome IV and VII. Both breakpoints occur in regions that have low divergence compared to the genome wide average (1 SNPs every 4.78 bp, red dashed line), consistent with local patterns of sequence similarity dictating the location of recombination breakpoints (Fig. 5E). Overall, the peculiar AIS genome structure is hard to explain with current models of *Saccharomyces* genome evolution. The formation of a full Sc x Sk hybrid with a sequential loss of 14 Sk chromosomes and re-diploidization of the corresponding Sc chromosomes or the partial transfer of two Sk chromosomes into a Sc strain represent two possible routes. The chromosomal heterogeneity of the AIS strain is likely responsible for the meiotic sterility, with gametes formed but fully unviable. Furthermore, a surprising and unique case of genetic code change resulting from a C to A mutation present on both haplotypes at position 35 in the original tR(ACG)K gene located on chromosome XI in the Sc part of the genome (between YKR026C and YKR027W), generated a novel tRNA^Arg^ with an AAG anticodon that would translate the CUU-Leu codons, and possibly the CUC-Leu codons as Arginine instead of Leucine. There are 5 other copies of the original tR(ACG) gene that most likely ensure the correct decoding of the CGT-Arg codons. The CUU-Leu codons are probably also normally translated as Leucine by the 3 original copies of the tL(UAG) genes but a certain level of ambiguous translation is expected from the presence of this novel tR(AAG) gene.

#### Dynamics of tDNAs multigene families

Multigene families of tDNAs are transcribed by the RNA polymerase III and serve as genomic targets for *de novo* transposition of the *Ty1* to *Ty4* elements (Bonnet and Lesage, 2021). As a result, tDNAs are usually located in complex repetitive regions of the chromosomes that cannot be assembled by short read genome sequencing. We identified 310 orthologous tDNA gene families by looking for tRNA gene copies in different strains that shared the same anticodon and were flanked by the same protein coding genes, at least on one side (Supp. Table 13). We used these families to look at dynamics of the tDNA gene repertoire in terms of codon modifications and gene family expansions and contractions. We focused on 100 haploid and homozygous diploid genomes (STAR METHODS, Supp. Fig. 22A and B), which presented comparable and consistent numbers of tDNAs, from 270 to 278 tRNA (with a mean and median of 275, Supp. Table 13). The tDNA repertoire is composed of 41 species of isoacceptors shared by all isolates. Two families underwent a mutation in the anticodon of one tDNA member. One mutation resulted in the formation of a new tRNA species in a single strain, CBK isolated from an insect in Germany. This tRNA arose from a G to T point mutation which transformed a tyrosine tDNA anticodon from GTA (tY(GUA), located on chromosome X) into a TTA nonsense suppressor (*SUP7*(UUA)). This *SUP7* gene has been previously detected in a clinical isolate, YJM421, where it contributed to a pair of Dobzhansky–Müller incompatibility by allowing it to read through the specific premature stop codon in *cox15^stop^* (Hou *et al*., 2015). Surprisingly, this premature stop codon is not present in the mitochondrial genome of CBK. The second family underwent an anticodon mutation in CEQ_1a, an African Palm wine isolate, where it resulted in a synonymous change from a tL(CAA) into tL(TAA) on chromosome 1 (Supp. Table 14).

We next investigated the plasticity of the tRNA gene repertoire by inventorying all gene gain and loss events (Supp. Table 13). We found that 248/310 families were conserved in all 100 isolates, while the others separated into two distinct categories depending on the number of strains they comprised. On the one hand, 35 tDNA families were found in less than 5 strains, suggesting that they were acquired by a recent tRNA gene gain event. On the other hand, 26 families were found in more than 90 strains suggesting that they underwent recent tRNA gene losses in 1 to 10 isolates (Supp. Fig. 22C). For 14 of them, the gene loss occurred in a single isolate. In the 10 remaining cases, the losses are not necessarily shared by closely related strains, suggesting that the same tDNAs can be independently lost several times in different isolates. In total, out of the 100 studied isolates, we found that 30 and 38 strains underwent 38 tDNA gene gains and 42 gene losses, respectively and all clades are affected by these events (Fig. 5F). Several strains accumulated several events, up to 5 in HN10 (BAM), isolated from rotten wood in China. Some clades preferentially accumulate one type of event, suggesting that functional constraints may favor the expansion or the contraction of the tDNAs gene repertoire. For instance, clades 9 (French Guiana human) and 10 (Mexican distillery/agave), plus the sister strain CQS_1a, underwent 7 tRNA gene gains but no loss. By opposition, the lab related clade (#13) underwent 5 losses and a single tRNA gene gain. Some tDNAs are recurrently lost, like the *tK(CUU)* which was lost 10 times independently but never gained (Fig. 5F, Supp. Table 14). Interestingly, tDNAs that were recently gained are located closer to the chromosomal ends than the conserved or lost tDNAs (Supp. Fig. 22D). There are 17/35 newly gained tDNAs that are located in the subtelomeres while none of the 248 conserved genes are and only 1/26 is subtelomeric among the lost ones. This suggests that the subtelomeres could serve as tRNA gene nursery where new copies are gained by segmental duplications associated with the junction of translocated segments from other chromosomes. The other 18/35 newly gained tDNAs that are located outside of the subtelomeric regions also result mainly from segmental duplications, either dispersed or in tandem.

#### The genealogy of Ty elements reveals their evolutionary dynamics

We inventoried all complete and truncated copies of retrotransposons and their solo LTRs from the five families of retrotransposons, *Ty1* to *Ty5* (Supp. Table 11) and found that Transposable Elements (TEs) are the other main contributors to genome size variations, along with the Y’ elements as described above (Supp. Fig. 18A and B). The second largest genome (12.65 Mb) is from a monosporic isolate (AMM_1a) derived from a homozygous diploid strain isolated from a leaf tree in Taiwan (SJ5L12, Supp. Fig. 18B). This genome has undergone a strong transposition activity (Bleykasten-Grosshans *et al*., 2021) with a total of 128 Ty elements (120 complete and 8 truncated) while the median number in the population is 14.5 (Supp. Fig. 23A and B). This activity involved *Ty1*, *Ty2* and *Ty4* elements with respectively 64, 53 and 26 copies. It is interesting to note though that some genetic factors, other than Y’ and Ty elements, can also significantly contribute to genome size variation given that two genomes with a similar number of Ty and Y’ elements can differ by more than 500 kb (Supp. Fig. 18C).

We also looked for the *Tsu4* element, from the related yeast species *S. uvarum*, because it has been identified once in a strain (245) isolated from a rum distillery in the West Indies, during a survey of 336 *S. cerevisiae* whole genome shotgun assemblies. The strain 245 only contains a single and nearly complete copy of *Tsu4* (Bergman, 2018). The authors proposed that this element would have been firstly horizontally transferred from *S. uvarum* into *S. paradoxus,* as revealed by the high similarity between *Tsu4* elements in these two species and a patchy distribution in *S. paradoxus,* and then secondarily transferred from *S. paradoxus* into *S. cerevisiae* (Bergman, 2018). We found the *Tsu4* element in a strain, CEY647 (CQS_1a), isolated from a bat in French Guiana. Surprisingly, this strain has 9 complete copies scattered on 7 different chromosomes, demonstrating that this element is functional in *S. cerevisiae*.

We found that the TE content is highly variable between isolates in terms of number and types of elements (Supp. Fig. 23A and B), in accordance with what has been previously described at the population level based on short reads sequencing data (Bleykasten-Grosshans *et al*., 2021). However, short-read sequencing precluded the accurate position of TE insertion sites within chromosomes. By opposition, the ScRAP provides the location of all insertion sites. As for tDNAs, the reconstruction of the Ty genealogy was achieved by identifying the insertion sites that are shared between several genomes. Insertion sites were considered to be shared between two genomes when two transposable sequences (complete or truncated elements or solo-LTR) were surrounded by the same orthologous protein coding genes as defined in the annotation files (at least on one side). In order to avoid the potential problem of heterozygous TE insertions, we restricted their analysis to the set of 100 haploid and homozygous genomes used above for the tDNA analysis. We identified 426 insertion sites (Supp. Table 15) that were plotted along the chromosomes of the reference genome (Fig. 5G). The repartition of these sites across the population shows a U-shaped distribution with 50% of them being shared by less than 15 strains and at the other end, 26% being shared by more than 90 strains (Supp. Fig. 23C). There is a good correlation between the number of strains sharing an insertion site and the number of solo-LTRs in these sites (r = 0.95, (Supp. Fig. 23D). In other words, the most conserved insertion sites are the most solo-LTR rich. For example, the 60 sites conserved in the 100 strains contain on average 84% LTRs and only 14% complete elements whereas the 86 insertion sites present in a single strain only contain only 52% LTRs and 47 % complete items. Therefore, solo-LTR remnants are shared by a much higher number of isolates (83.75 on average) than complete elements (13.75 on average, Fig. 5G), suggesting that inter-LTR recombination is common. In line with this observation, none of the complete elements are shared by all 100 isolates, the most conserved complete elements being present in only 62 strains (a *Ty4* element on chrXIV, Fig. 5G, Supp. Table 15). Another manifestation of the poor conservation of complete elements at the population level is that 118/426 insertion sites contain no full-length copy at all (Fig. 5G). The 4 closely related strains from the Malaysian clade (BMB, BMC_2a, UWOPS034614 and UWOPS052272) contain average numbers of solo-LTRs (about 390) and truncated Ty copies (between 6 and 9) but are completely devoid of full-length element, suggesting that all functional copies were lost by recombination between LTRs in this clade. Interestingly, it was previously reported that the genome of the UWOPS034614 Malaysian strain underwent a massive rearrangement with 8 translocations resulting from four reciprocal translocations between 5 chromosomes as well as 4 subtelomeric translocations and a few intrachromosomal inversions that occurred in breakpoint regions containing tDNAs and Ty elements (Marie-Nelly *et al*., 2014; Yue *et al*., 2017). Here we found that the 4 Malaysian strains are among the most rearranged genomes of the ScRAP, with 14 translocations and between 6 and 8 inversions per genome (Supp. Table 6), in accordance with previous results. Increased genome plasticity in the Malaysian clade could therefore result from increased ectopic recombination between dispersed repeats in these genomes.

The frequent loss of complete elements by inter-LTR recombination at the population level is likely counterbalanced by an active process of *de novo* transposition. There are 61 sites in which only complete elements are found and never soloLTR, which strongly suggests that these sites correspond to recent insertions. In accordance with this hypothesis, the ⅔ of them (41/61) are found in a single isolate (Supp. Table 15). The other 20 insertion sites are shared between a small number of strains (between 2 and 7) and these strains are phylogenetic neighbors (Supp. Fig. 24). This is particularly visible in clade 13. Lab-related in which we see 30 new insertions in 7 sister strains, resulting from 5 and 1 recent independent insertion events of *Ty1* and *Ty2*, respectively(Supp. Table 15, Supp. Fig. 24).

In conclusion, the genealogy of the TEs revealed that the oldest sites that are shared by a greater number of strains, have undergone frequent inter-LTR recombination events which results in the frequent elimination of complete elements of the genomes. However, these complete elements actively transpose into new regions which are therefore less shared at the population level but comprise a higher proportion of complete elements. As a result, the chromosomal localization of complete elements is only poorly conserved at the population level unlike that of solo-LTRs.

## DISCUSSION

Understanding the genetic basis of simple and complex traits has been hampered by several factors, including the lack of complete knowledge of the causative genetic polymorphisms especially with respect to large-scale SVs. Using telomere-to-telomere *de novo* assemblies of a large panel of *S. cerevisiae* strains we predicted the whole structural diversity to contain approximately 6,000 events(Fig. 2B) of which we were able to characterise about 80%. We estimated that accessing the missing ∼1,200 events would require an additional ∼360 strains. We showed that SVs, notably insertions, deletions and duplications can impact the expression of genes located nearby. Additionally, we found that SVs have a strong potential to increase the diversity of the gene repertoire through gene truncations, duplications and fusions, calling for a redefinition of the PanORFeome at the species level and the exploration of the phenotypic impact of accessory genes (McCarthy and Fitzpatrick, 2019). The true contribution of both SVs and accessory genes to the missing heritability remains to be quantified but the ScRAP represents a critical genomic resource towards this goal. High-throughput approaches such as GWAS, multiplexed genome editing and deep mutational scanning can now be envisioned to determine the impact of SVs and accessory genes on phenotypic diversity and adaptation. Identifying causal variants beyond the SNVs, notably in subtelomeric regions which have remained largely excluded from most analyzes despite being enriched in SVs and accessory genes (Verstrepen and Klis, 2006), can also help to select new strains for industrial applications.

We found a median of 240 SVs per genome which represents an average density of 1 SV every 50 kb. By comparison, each human genome would contain >20,000 SVs (Ho *et al*., 2019), which corresponds to approximately 1 SV/150 kb, *i.e.* 3 times lower than in *S. cerevisiae.* In other eukaryotes that benefit from pangenome data, the SV density scales from 1 SV/90 kb in *Drosophila* (Chakraborty *et al*., 2019), 1 SV/38 kb in soybean (Liu *et al*., 2020), 1 SV/17 kb in rice (Qin *et al*., 2021) and up to 1 SV/4 kb in silkworm (Tong *et al*., 2022). We also found a clear positive correlation between the numbers of SVs and SNVs/INDELs accumulating within genomes. A genomic clock that would synchronize protein divergence and chromosomal rearrangements in bacteria and yeast (Vakirlis *et al*., 2016; Puigbò *et al*., 2014), however this clock seems to tick at a different pace depending on the ploidy and zygosity of the genome. SVs seem indeed to preferentially accumulate in heterozygous and higher ploidy genomes (Fig. 3B). One possibility would be that SVs would be better tolerated in higher ploidy genomes as their deleterious effects (gene deletion or disruption, dosage imbalance, etc) could be more efficiently buffered. Alternatively, we cannot exclude that the rate of SV formation would increase with ploidy, as was already suggested for aneuploidies (Gilchrist and Stelkens, 2019).

We characterized a novel type of genetic diversity in the form of complex aneuploidies where aneuploid chromosomes are associated with interchromosomal translocations and/or large deletions. Yet, what we define as complex aneuploidies is similar to previously reported cases of ‘segmental aneuploidies’ in which two copies of the left arm of chromosome 5 were fused around a single copy of the centromere in several azole resistant strains of *Candida albicans* (Selmecki *et al*., 2006). However, in many cases segmental aneuploidies contrast with our definition of complex aneuploidies whereby a centromere must be contained within the copy-number modified region. For instance, in humans, this is the case for chromosome arm aneuploidies (CAAs) which occur frequently in cancer and even more frequently than whole chromosome aneuploidies in certain cancer types (Shukla *et al*., 2020; Taylor *et al*., 2018). These CAAs have the potential to represent complex aneuploidies as we have described here. However, due to the complex repetitiveness of human centromere regions, only recently have these regions been reliably assembled (Altemose *et al*., 2022) and therefore centromere complexity may have hidden these cases in other aneuploidy rich data such as human cancer. Several studies in *Saccharomyces* reported a significant negative correlation between chromosome size and the rate of simple aneuploidy (Gilchrist and Stelkens, 2019), probably because the fitness cost of extra chromosomes is proportional to the total number of genes present in the excess chromosomes. In line with this, CAAs frequency in human cancers is inversely related to arm length (Beroukhim *et al*., 2010). On the other hand, we showed that larger chromosomes are more often involved in complex aneuploidies than the smaller ones. Therefore, we hypothesize that complex aneuploidy offers a powerful adaptive route that would be inaccessible to simple aneuploidies by allowing to combine the benefits of increased copy number of selected genes from large chromosomes and reducing the cost of gene imbalance through either large deletions that encompass deleterious genes or translocations. In addition, they are widespread in the population, as the re-analysis of the genomes of 993 strains showed that up to 18% of all aneuploidies could be complex, while they should be extremely rare as they are expected to occur at a rate corresponding to the product of the rates of aneuploidy and SV formation. Therefore, complex aneuploidies could possibly be adaptive and selected for in response to harsh environments, as was demonstrated for simple aneuploidies (Mulla *et al*., 2014). Further work is needed to characterize the molecular mechanisms by which complex aneuploidies are formed and to determine their specific adaptive potential.

In the near future the objective will be to generate an unified, complete and accurate representation of the genomic diversity at the species level. The ScRAP provides a solid foundation for this purpose and will drive the transition from the use of a single linear reference genome to that of a pangenome devoid of reference bias, which, as the ScRAP contains, requires the integration of all ploidy levels, including polyploid, all simple and complex aneuploidies and all heterozygous variants within fully phased and contiguous haplotypes. This will enlighten our knowledge on genome evolution as exemplified in this work where, in addition to the characterization of the SV landscape, we report for the first time the individual telomere length variation across the entire population, the explanation of how large HGT regions colonize telomeres and can subsequently re-localise in chromosome core, the description of a novel type of chromosome-scale admixture and the reconstruction of the evolutionary dynamics of tRNA multigene families and Ty elements.

What will be the best data structure to efficiently describe and interrogate a large and growing collection of genomes within the pangenome? This question is at the heart of an active field of research whose objective is to develop new bioinformatic methods required to produce a graphical pangenomic reference system (Eizenga *et al*., 2020). The ScRAP provides an exceptional dataset that will be instrumental for the development of these tools and the emergence of population pangenomics.

## Acknowledgments

We thank B. Dujon and B. Llorente for their valuable feedback on the manuscript. This work was supported by the Agence Nationale de la Recherche ANR-16-CE 12-0019 and ANR-18-CE12-0004. This work was also partially supported by the ANR-15-IDEX-01 (to G.L.), Fondation pour la Recherche Médicale (EQU202003010413 to G.L.), the European Research Council (ERC Consolidator Grant 772505 to J.S.), the Guangdong Basic and Applied Basic Research Foundation (2019A1515110762 to J.-X. Y.), Guangdong Pearl River Talents Program (2019QN01Y183 to J.-X. Y.), and National Natural Science Foundation of China (32070592 to J.-X. Y.). J.S. is a Fellow of the University of Strasbourg Institute for Advanced Study (USIAS) and a member of the Institut Universitaire de France.

## STAR□METHODS

### ScRAP strain selection

The rationale for the selection of the 100 strains *de novo* sequenced in this study was based mainly on the knowledge gained from the 1,011 genome project (Peter *et al*., 2018; De Chiara *et al*., 2022). We selected one per clade and subclades, with a good sporulator phenotype. We selected some strains with a known signature of SV (e.g. AIF with segmental duplications). The AIS strain that contained chromosome-scale introgression was first detected in the 1,011 work but excluded because of its complex genome structure. We also selected strains known to carry large HGT events. The 31 diploids (10 nearly homozygous and 21 highly heterozygous) that were unable to sporulate or to produce viable spores were sequenced in their original ploidies. Note that, as an exception, BAF was sequenced as a diploid despite the fact that it is sporulating well and has a good spore viability.

### DNA extraction and sequencing

#### ONT library preparation and sequencing

We grow yeast cells in 10 - 15 ml YPD at 30°C overnight (220 rpm). A total number of cells less than 7 x 109 were used for DNA extraction. High molecular weight (HMW) DNA was extracted by QIAGEN Genomic-tip 100/G according to the “QIAGEN Genomic DNA handbook” for Yeast. DNA quantity and length were controlled by Qubit dsDNA HS Assay and PFGE respectively. Library preparation and ONT sequencing were performed based on the protocol of ‘1D Native barcoding genomic DNA with EXP-NBD104 and SQK-LSK108’ when using FLO-MIN106 MinION flow cells and protocol of ‘1D Genomic DNA by Ligation with EXP-NBD104 and SQK-LSK109 - PromethION’ when using the V2 FLO-PRO002 flow cell. These protocols are available from Oxford Nanopore Technologies Community.

For sequencing library preparation, up to 2 ug of HMW DNA per sample was used to start library preparation. DNA repair and end preparation were performed using the NEBNext FFPE DNA Repair Mix with the following reaction setup: 48 ul DNA, 3.5 ul NEBNext FFPE DNA Repair Buffer, 2 ul NEBNext FFPE DNA Repair Mix, 3.5 ul Ultra II End Prep Reaction Buffer, and 3 ul Ultra II End Prep Enzyme Mix; 20°C for 15 min followed by 65°C for 15 min. Afterward, the DNA size selection was carried out using AMPure XP Beads (1:1 ratio) followed by native barcode ligation (22.5 ul DNA, 2.5 ul native barcode provided by EXP-NBD104 kit and 25 ul Blunt/TA Ligase Master Mix; 25°C for 20 min). After another round of AMPure XP bead clean-up (1:1 ratio), the samples were pooled together and adaptors were ligated for the pooled sample at 25°C for 15 min (65 ul DNA, 5 ul AMII provided by EXP-NBD104 kit, 20 ul NEBNext Quick Ligation Reaction Buffer, and 10 ul Quick T4 DNA Ligase; 25°C for 15 min). The adaptor-ligated DNA was cleaned up by adding a 0.4× volume of AMPure XP beads followed by incubation for 5 min at room temperature. When using SQK-LSK108 kit for FLO-MIN106 MinION flow cells, twice 140 ul ABB washes were performed. When using SQK-LSK109 kit for FLO-PRO002 flow cells, twice 250 ul L Fragment Buffer (LFB) washes were performed. The final library was eluted in 15 ul Elution Buffer and loaded into the MinION or PromethION flow cells according to the ONT manuals. Raw fast5 files were basecalled using Guppy (v3.4.5) followed by Porechop (github.com/rrwick/Porechop) removal of adapters and barcodes. The entire project generated close to 204 Gbp of Nanopore sequencing data. Sequencing statistics are detailed in Supp. Table 16. For fast5 file storage/sharing, single-fast5 files were stripped of basecalling data using Picopore (github.com/scottgigante/picopore), ensuring all files contain only data necessary for re-basecalling. Next single-fast5 files were converted into multi-fast5 files using the ont-fast5-api (github.com/nanoporetech/ont_fast5_api) *single_to_multi* command, followed by the *fast5_subset* command to generate strain specific fast5 files containing fast5 files for all reads within each strain specific fastq file. This was done to reduce the complexity of reanalysis using fast5 files from strains run with both/either multiple barcodes and across several flowcells and remove fast5 files for reads of insufficient quality. All adapter/barcode free fastq files and their associated strain-specific fast5 files are available under the accession PRJEB50706/ERP135326.

#### Illumina sequencing

We grew yeast cells cultures overnight at 30° C in 20 ml of YPD medium until early stationary phase. We harvested cells by centrifugation and extracted total genomic DNA using the QIAGEN Genomic-tip 100/G according to the manufacturer’s instructions. Genomic Illumina sequencing libraries were prepared with a mean insert size of 280 bp and subjected to paired-end sequencing (2 x 100 bp) on Illumina HiSeq 2500 sequencers. All paired-end Illumina reads are available under the accession PRJEB50706/ERP135326.

### Genome assembly pipelines

All processes are illustrated in Supp. Fig. 25.

#### Nuclear chromosomes

Raw fastq files were treated with porechop (v0.2.3) to remove both adapters and barcodes (github.com/rrwick/Porechop), downsampled with Filtlong (v0.2) for a maximum of 40X per strain (github.com/rrwick/Filtlong) then assembled by both Canu (v2.0) (Koren *et al*., 2017) and SMARTdenovo (https://github.com/ruanjue/smartdenovo). This generated two assemblies per strain that were treated separately. The contigs were polished using the same fastq data used for assembly. First both the Canu and Smartdenovo assemblies are polished by Racon (v1.4.3) for one and three rounds respectively (Vaser *et al*., 2017). This is followed by two rounds of Medaka (v1.2.2) (github.com/nanoporetech/medaka). The Nanopore polished contigs were then polished using Illumina data with Pilon (v1.22) for three rounds (Walker *et al*., 2014). The hybrid-polished contigs were scaffolded against the reference genome using Ragout (v2)(Kolmogorov *et al*., 2014), and then manually curated to correct for un-scaffolded or badly scaffolded contigs, to generate a haploid assembly. Potential haplotigs and mitochondrial contigs were then removed based on a reference-based renaming of contigs/scaffolds. Assemblies were descaffolded and manually inspected for negative gaps using Gap5’s ‘Find internal joins’ option (v1.2.14)(Bonfield and Whitwham, 2010). Contig overlaps were considered negative gaps if the same two contigs were previously scaffolded in this gap and if the overlap was greater than 10kb. In these cases, the negative gaps were closed by generating an overlap consensus. Those assemblies that had negative gaps removed were then re-polished by Pilon once to correct for any errors introduced during the consensus generation step. Finally, assemblies were then re-scaffolded with Ragout followed by manual curation, generating the final assemblies.

For all haploid/homozygous and haplotype-collapsed assemblies, the two alternate versions of each assembly (Canu and SMARTdenovo) allowed us to evaluate possible misassemblies or assembly artifacts by aligning them against the S288C reference genome as well as against one another using MUMmer’s nucmer/mummerplot tools (v4)(Marçais *et al*., 2018) and manually inspecting for structural differences. In the case where a structural discrepancy was found between the Canu and SMARTdenovo versions, the best assembly corresponded to the one with the reference-like structure. Indeed, both assemblies are *de-novo* structures and therefore an assembly sharing a reference-like chromosome structure is more likely to be real than the other uniquely rearranged assembly. Discrepancies in both assemblies were present only in heterozygous strains, with the exception of three homozygous/haploid strains (BAM, BGP and CRB). Due to this, the ‘best’ assembly for BAM and BGP contains a combination of contigs from Canu and SMARTdenovo (BGP contains all but chromosome V from the SMARTdenovo assembly and BAM contains primarily Canu except for chrXII, chrIII and chrVII_IV from SMARTdenovo). The CRB discrepancy was unresolved and therefore the choice of the best assembly relied on the assembly statistics below. In the absence of assembly structure discrepancies, assemblies were compared using basic genome statistics and those generated by MUMmer’s dnadiff tool (Marçais *et al*., 2018). We used the following values for comparison; number of scaffolds, number of contigs, percentage of reference covered, percentage of reference identity and genome size, in order of decreasing weight. The aim was to select a genome with an ideal number of 16 contigs/scaffolds and maximize the remaining values. Contig count was reduced by one if a scaffold was present within the rDNA array on chrXII. Selecting the ‘best’ phased assembly for polyploid strains relied purely on stats due to regular structural discrepancies between assemblies. Diploid heterozygous strains also contained structural discrepancies, however, the ‘best’ assembly relied upon phasing statistics (see Haplotype phasing).

#### Mitochondrial chromosomes

The mitochondrial de novo genome assemblies were constructed from the Illumina paired-end reads with a pipeline derived from (Tao *et al*., 2019). Short reads were down-sampled to sets of 500,000, 600,000, 700,000 and 800,000 paired-end reads with seqtk (https://github.com/lh3/seqtk). Each dataset was de novo assembled with A5-miseq (Coil *et al*., 2015) and mitochondrial contigs were identified through similarity searches to the *S. cerevisiae* mitochondrial reference sequence (accession number KP263414.1). For each strain, a representative assembly was selected based on the number of contigs and their length. The one-contig assemblies were subjected to circlator for circularization (Hunt *et al*., 2015). For the non-circularisable ones, long reads were screened and used for manual circularisation if possible. A custom python script was used to set the starting position of the sequence to the ATP6 gene initiation codon.

### Haplotype phasing

All processes are illustrated in the Supp. Fig. 25. In order to test the following pipeline a test dataset was generated from a strain, SO002, generated in the lab by crossing two stable haploids, YLF161 and YLF132 (Louvel *et al*., 2014), from West African (DVBPG6044) and North American (YPS128) backgrounds respectively. After generating the diploid, it was then sequenced by nanopore and illumina as with other strains within this study. This dataset was created to remove biases from generating diploids artificially through in silico read subset merging and as both parental strains have reference quality assemblies for comparison (Yue *et al*., 2017). The results of the pipeline, the analysis, the results and raw data can be found at (https://github.com/SAMtoBAM/PhasedDiploidGenomeAssemblyPipeline).

For the 21 diploid heterozygous strains, the following phasing pipeline was performed three times per strain, with each run only changing the genome backbone used for read alignment. The three backbone genomes used were the S288c reference genome and both canu and SMARTdenovo collapsed de-novo assemblies. Illumina reads were aligned to the genome assembly using BWA-MEM (v0.7.17) (REF), converted to bam and sorted with samtools sort (v1.11) (REF). Next, variants were called using GATK MarkDuplicatesSpark and HaplotypeCaller (v4.1.8.1) (REF) and filtered for a minimum mean DP of 30 with vcftools (v0.1.5) (REF). ONT reads were aligned to each assembly using Minimap2 (v2.13) (REF), converted to bam and sorted with samtools sort (v1.11). The resulting short-read based VCF and LR based BAM were given as input to Whatshap’s phase tool (v1) (REF) to phase the variants and produce a phased VCF. The phased VCF and LR based BAM were provided as input to Whatshap’s haplotag tool to phase the reads and produce a phased BAM. Reads are then split by the Haplotype tag (HP1/HP2) and combined with unphased reads to generate one set of reads for each haplotype.

Phasing statistics were evaluated in order to determine the best assembly for phasing and in turn the best de novo assembly for each strain. In order to evaluate phasing contiguity new phasing statistics were created, the V90 and the nV90. The V90, inspired by the L90 and N90, is the minimum number of phased blocks to cover 90% of all phased variants. The nV90 is the normalized V90, dividing the V90 by the total number of contigs in which the V90 blocks are present. Therefore, the ideal scenario is that each block corresponds to a unique contig giving an nV90 of 1. The highest nV90 was used to select the ‘best’ assembly. In 3 cases where the nV90 were equal in both Sdn and Canu assemblies, the assembly with the greatest total sum of phased basepairs was chosen. First, comparing the use of the de-novo assemblies versus the reference, identified that de-novo assemblies consistently performed better (Supp. Fig. 26). Secondly, out of the two *de novo* assemblies, the phasing results determined the ‘best’ assembly and the phased reads were used downstream for haplotype assembly. 6/21 genomes chosen based on phasing stats disagreed with the assembly choice based on stats alone.

For polyploid strains, we used the nPhase tool with default parameters to phase each polyploid using both long and short reads (Abou Saada *et al*., 2021). Once we obtained raw results using the nPhase pipeline command, we ran the nPhase cleaning command using default parameters to improve contiguity and eliminate short, uninformative haplotigs.

### Genome annotation

All processes are illustrated in the Supp. Fig. 25.

All nuclear and mitochondrial assemblies were annotated with the LRSDAY pipeline (v.1.6.0) (Yue and Liti, 2018). Based on sequence homology comparison, LRSDAY automatically identifies protein-coding genes, centromeres, transposable elements, telomere associated X’- and Y-elements as well as important mitochondrial RNAs. Three types of transposable elements were annotated: complete and truncated Ty, as well as solo LTR, each of them being classified in Ty1, Ty2 (or Ty1/Ty2 for solo LTR), Ty3, Ty4 and Ty5 families. For each input assembly, the annotation of different genomic features were combined into a single GFF3 file, which were further examined, resorted, and verified by GFF3toolkit (v2.1.0). Those annotated protein coding genes with disrupted ORFs were labeled as pseudogenes during this process.

### Analysis of complex regions

#### Subtelomere dynamics

The subtelomeric regions were annotated and named in the same way as our previous study proposed (Yue *et al*., 2017). Manual examination and adjustment were further applied to curate subtelomeric ends with incomplete sequence information or substantial reshuffling (Supp. Table 17).

#### tDNAs and Ty elements

We defined a consistent set of 100 haploid or homozygous genome assemblies for the analyses of the dynamics of tDNAs and Ty elements by first excluding the diploid, triploid and tetraploid phased assemblies because they contained numbers of annotated tDNA and Ty copies that were proportional to their ploidy and therefore difficult to compare with haploid and collapsed genome assemblies (Supp. Fig. 17, Supp. Table 11). We also removed 8 haploid genomes from one specific study (Bendixsen *et al*., 2021) because they contained a much lower number of tDNAs than all other genomes of the dataset (Supp. Fig. 22A), probably indicative of local assembly errors. We finally excluded collapsed assemblies from heterozygous genomes because they showed some discrepancies with their cognate phased assemblies, suggesting possible assembly problems in these complex regions and also because they contained the most extreme numbers of tDNAs per genome (Supp. Fig. 22B).

### Telomere detection

We developed a python package called *Telofinder* to determine the chromosomal location and size of telomere sequences in yeast genomes assemblies (https://telofinder.readthedocs.io/en/latest/). Telomere detection is based on calculation of both the DNA sequence entropy and the proportions of the “CC”, “CA” and “AC” dinucleotides in a 20 bp sliding window. *Telofinder* outputs two csv and two bed files containing the telomere calls and their coordinates, either as raw output or after merging consecutive calls. We run *Telofinder* on all 394 *de novo* and 24 previously available nuclear genome assemblies with the -s option set to -1 to scan entire genome sequences.

### Aneuploidy detection

#### Aneuploidy detection in the ScRAP

Illumina reads were aligned to the reference genome using bwa-mem and coverage calculated using bedtools genomecov (-d -ibam). Genome coverage was then visualized for each strain separately with centromere positions. Aneuploidies were then manually annotated. In order to validate complex aneuploidy structures, we used additionally both nanopore reads and genome assemblies. Nanopore reads were aligned to the reference using minimap2 and visualized using tablet (Milne *et al*., 2016). Raw and finalized genome assemblies were aligned with nucmer against the S288c reference assembly and other assemblies and visualized as a dotplot. Additionally, by analyzing raw genome assemblies, four complex aneuploidy chromosomes which were completely or partially assembled, at least containing the complex region, were identified and extracted (CBS1586/AHG +1*chr10c; CBS457/AIF +1*chr11c; CBS4255/ASB +2*chr9c; CBS1489/ASG +1*chr3c).

#### Aneuploidy detection in a dataset of 909 strains

The aneuploidy detection pipeline using (Peter *et al*., 2018, 011) and (Strope *et al*., 2015) illumina data is available at https://github.com/SAMtoBAM/aneuploidy_detection and consist of the following steps:

1. Illumina reads were aligned with bwa mem and the coverage calculated using bedtools *genomecov* (-d -ibam) .
2. Coverage was binned the median coverage in 30kb bins with a 10kb slide using both bedtools *makewindows* and *map* and remove regions covering 15kb (half a window) from each chromosome end (to reduce telomere mapping/variation issues).
3. Coverage was normalized to the genome wide median and regions extracted if they deviated by +-0.7*(1/n). 0.7 gives some leniency to coverage deviation considered sufficient for a change in copy number.
4. Deviating bins were aggregated, allowing for a gap of <=10kb (the size of 1 slide).
5. Aggregated bins were split into two types depending on whether they overlapped a centromere or not, called centromere related (CR) and non-centromere related (NCR) respectively.
6. The size of CR regions was increased to the sum of all regions within the same chromosome with the same deviation e.g. +1, +2, -1 etc. to give a CR-sum. We then removed any CR-sum < 50kb and calculated the difference between this CR-sum and the chromosome size (minus the 30kb removed from the ends).
7. All CR-sums with a difference in size >100kb (i.e. a the CR does not cover region/s summing to 100 kb or more) were labeled as complex and the rest as simple
8. Plot the normalized coverage of all aneuploidies and manually curate the list in order to remove false positives and adjust the complex-simple classification. During the manual curation, 34 complex stayed complex, 347 simple remained simple, 4 complex were removed as false positive aneuploidies (due to the impact of the ‘smiley-effect’), 32 moved from complex to simple, 10 moved from simple to complex. The move from simple to complex was always due to the threshold of size (the size difference of these ten CRs was 96, 90, 82, 80, 72, 72, 66, 66, 50 and 40kb). All examples of this reclassification from simple to complex were distinct.
9. Identify any NCRs >100kb which are present within a strain containing an aneuploidy detected above. Label as complex aneuploidy-related and use in the less conservative estimate of complex aneuploidy count.

The newly generated large aneuploidy dataset overlaps, by strain and chromosome, 88% (303/343) with that of the (Peter *et al*., 2018) dataset. This leaves only 40 aneuploidies (12%) not re-detected. Of this 40, manual inspection identified that 9 were clearly false positives in (Peter *et al*., 2018), 8 came from the same strain and are an issue of defining whether 8 chromosomes were lost or the other 8 were gained, 1 contains a large increase in coverage close to the centromere but not covering and 3 contain a slight change in coverage but are well below the set threshold for indicating a change in copy number. Therefore, only 19 missed aneuploidies are likely true false negatives in the new dataset. The overlap between datasets additionally shows 120 previously undetected aneuploidies in the new dataset. Of these 120, 35/120 (29%) are complex aneuploidies, as compared to 9/303 (2%) found within the overlap.

### SV detection

All processes are illustrated in the Supp. Fig. 25.

MUM&Co (v3.8) (O’Donnell and Fischer, 2020) called SVs separately on all genomes generated, including both Canu and SMARTdenovo, both phased and collapsed, 4 complex aneuploid chromosome assemblies and 23 public assemblies, totalling 3229 assemblies. For phased polyploid genomes, each individual phased block assembly was run separately and the -ml/--minlen was increased to 100. All raw calls were aggregated and then both insertion events involving scaffold fragments (N’s) and calls within the region surrounding the rDNA cluster on chrXII in the reference (position 450-490kb) were removed. This gave a total of 95893 raw, unvalidated calls. For any call to be validated, and aggregated into a non-redundant call, it had to match the following criteria with another call of the same type in another assembly: for Deletions, Duplications and Contractions events: matching start and end reference position +-300bp; for Insertion events: matching start and end reference position +-300bp and Size of inserted fragment (+-10%); for Inversion events: matching start and end reference position +-6.5kb; for translocation events: matching start and/or end reference position +-6.5kb and matching chr involved at border and matching position within chr involved at border +-6.5kb.

If any calls did not match another call using these rules, it was removed. Therefore, validation requires a call to be found within at least two genome assemblies, whether within or between strains. An exception is made for both duplications and contractions which can be validated within a single genome if the region contains more or less than one additional copy, i.e. triplicated, and therefore contains multiple duplication calls for a single region. However, there was only one occurrence of a duplication being validated within only a single assembly. After the aggregation of calls into a single non-redundant event, an average value for the positions and size was taken. Furthermore, the orientation for translocation events (following border events from VCF formatting, e.g. [chrI:1[) was taken by consensus. In the case that there was an equal number of both orientations, either contigs were reverse complemented followed by re-calling SVs (commonly found in the case of polyploid phased assemblies that were not reference orientated prior to SV calling) or calls were split in the case of likely over-aggregation. Additionally, due to each phased block for polyploid strains having been analyzed for SVs individually, the discontinuity of each ‘assembly’ added large false positive deletions. Due to this, all large deletions up to a size of 20kb were removed if not manually verified by coverage analysis, as performed for aneuploidy detection (refer to Aneuploidy detection). Using these rules, 91645 (95.6%) were considered validated, which when split by SV type ranged from 78-99% for translocations and contractions, respectively (Supp. Fig. 27). Additionally, an average of 95.7% of all variants within each strain were validated. Notably however, genomes that were not assembled within the frame of this project had a lower average within-strain validation due to only containing a single assembly per strain. Furthermore, many from Bendixsen et al. (2021) have genetically diverse East Asian origins and therefore reducing the likelihood of sharing an SV with another strain within this cohort (Supp. Fig. 28). The total non-redundant dataset contains 4809 SVs.

For diploid genomes, validated SVs were considered homozygous if found in at least one genome from both HP1 and HP2 for the same strain. Alternatively, SVs were considered heterozygous if only a single haplotype (HP1 or HP2), from the same strain, were found to contain the SV. In this respect, there is an assumption that the phased genome contains two complete haplotypes due to the fact that unphased reads were also added to phased diploid genome assemblies. In contrast, polyploid phased genomes contain only phased reads and therefore the number of haplotypes can vary for any reference position. Therefore, in order to label calls homozygous or heterozygous, first the number of haplotypes must be estimated. Polyploid phased genomes were first aligned to the reference using Minimap2 (using options: -ax asm5 --secondary=no) and coverage per reference base pair calculated. The number of haplotypes in regions of deletions and contractions was determined by the median coverage 20kb up and downstream of the event. For all other events, the number of haplotypes was calculated using the median coverage of both 20kb up and downstream and within the event itself. All SVs that were present within fewer phased assembly blocks than the number of predicted haplotypes were considered heterozygous. Lastly, in regions where the number of predicted haplotypes was greater than the ploidy +2 (e.g. 5 in a triploid), events were considered homozygous.

Insertions, deletions, duplications and inversions can be further broken down into whether they involve novel or repetitive regions of the genome. To calculate the material involved in each SV was searched using BLAST against the reference. In order for an insertion to be novel, the material should not already be present. However, for deletions, duplications and inversions, they by definition are already present and therefore are considered novel if only present once. Therefore, after the BLAST search, matches were filtered to have an e-value <0.001 and a base identity >90%. The sum of the remaining lengths was then calculated. This sum needed to be above 50% or 150% of the query length to be considered a repetitive region in insertions and deletion/duplications respectively.

One way to look at gene impact is to categorize the type of impact. Four categories have been created, ignorant of the resulting ORF(s) and its/their direction: (i) Contained = At least one entire gene is contained within the SV (excludes insertions and translocations), (ii) Disruption = A single gene is present at the border of an SV (excludes insertions), (iii) Within = the entirety of the rearrangement is confined within a single gene (excludes translocations) and (iv) Fusion = Both borders of an SV interact with different genes which would therefore bring them into frame in a way (excludes insertions). The categories are not mutually exclusive as a single event can both contain entire genes and disrupt/fuse others at the border. However, an event cannot disrupt and fuse and/or be within. Additionally, the gene repertoire can be filtered to only contain essential genes and the analysis repeated. However, fusion can happen between essential and non-essential genes. Therefore, a 5th category in a way was added: (v) Fusion-between-essential-and-nonessential = Both borders of an SV interact with different genes which would therefore bring them into frame in a way (excludes insertions). One of those genes is an essential gene.

### SVs and gene expression impact

#### Determination of SV-gene pairs

SV-gene pairs were evaluated using bedtools. For SVs overlapping CDS, bedtools intersect was used to identify the pairs. A supplementary awk filter was applied to specifically identify CDS fully located within SVs. For SVs within intergenic regions, bedtools closest was used to identify the pairs, either by identifying the two SVs closest to a CDS in the case of indels (using the –io and –id or –iu options) or by identifying the CDS closest to each boundary of an SV, as well as the ones that might overlap SV boundaries. IN the case of inversion events, only the pairs involving the genes closest or overlapping the associated inversion breakpoints were investigated. Additionally, in the case of indels, SVs were associated with a CDS only if they were located in the intergenic space between observed CDS and the next, both up- and downstream.

#### Evaluation of the impact of SVs on the expression of neighboring genes

For each of the SV-gene pairs obtained at the previous step, the 51 studied strains were split into two groups: strains with and without the SV. Then, the expression values of the studied gene in each of the strains were ranked and normalized between 0 and 1, and then used to evaluate differential expression by performing a two-sided Wilcoxon-Mann-Whitney test between the with- and without-SV groups. This statistical analysis was performed using R, version 3.5.1.

### Phylogenetic reconstruction

#### Ortholog-based nuclear tree

For the nuclear genome, the proteome sequences of 181 input genomes (with 23 outgroups *Saccharomyces* species) were used for phylogenetic analysis. A total of 1612 one-to-one nuclear ortholog groups were identified by Proteinortho (version: 6.0.25; options: --check -selfblast -singles). For each ortholog group, the protein and CDS alignment of were generated by MACSE (version: 2.04; options: -prog alignSequences -gc_def 1 -seq $i.species_relabeled.fa -out_NT $i.macse_NT.aln.fa - out_AA $i.macse_AA.aln.fa and -prog exportAlignment -align $i.macse_NT.aln.fa - codonForFinalStop --- -codonForInternalStop NNN -codonForInternalFS NNN - codonForExternalFS --- -charForRemainingFS - -out_NT $i.macse_NT.aln.tidy.fa - out_AA $i.macse_AA.aln.tidy.fa). A concatenated super matrix of the 1612 ortholog-based CDS alignment was further generated with different partitions defined corresponding to different ortholog groups. This super matrix and its associated partition definition were used for maximal likelihood tree building by IQtree (version 1.6.12; options: -spp $prefix.concatenated.cds.partition.txt -s $prefix.concatenated.cds.tidy.fa -m MFP -bb 1000 -alrt 1000 -nt $threads -pre $prefix.iqtree -safe). 1000 rounds of ultrafast bootstrap (UB) and approximate likelihood-ratio test (aLRT) were used to assess the branch supports.

#### SNP-based nuclear tree

The input vcf file (matrixSam.snp.vcf.gz) was generated using GATK4’s HaplotypeCaller and GenotypeGVCFs (v4.1.4.1) with bwa-mem (v1) aligned illumina reads. The resulting multi-sample VCF was then filtered for variants with a Quality-by-Depth (QD), StrandOddsRatio (SOR), FisherStrand (FS), Mapping Quality (MQ), MappingQualityRankSum (MQRankSum) and/or ReadPosRankSum (RPRS) of more than two standard deviations from the average. Lastly, variants were removed from regions labeled as repetitive by RepeatMasker and/or in (Jubin *et al*., 2014) to generate the final VCF.

We used the vcf2phylip python script (https://github.com/edgardomortiz/vcf2phylip) (versions: 2.8; options: -I $input_vcf –resolve-IUPAC -o S288C –fasta –output-prefix) to convert the vcf file into the fasta format. The corresponding SGDref entry in fasta format was extracted based on the reference allele column of the input vcf file. MAFFT (version: 7.471; options:--auto --thread $threads --preservecase -- addfragments) was used to align these two fasta files by using the extracted SGDref entry as the reference sequence for alignment. The resulting alignment was further filtered by ClipKIT (version: Github commit cccc8bf; options: -m gappy). The filtered alignment was fed into IQtree for tree building (version 1.6.12; options: -s $prefix.fasta -m GTR+ASC -bb 1000 -alrt 1000 -nt $threads -pre $prefix.iqtree -safe). A thousand rounds of ultrafast bootstrap (UB) and approximate likelihood-ratio test (aLRT) were used to assess the branch supports.

#### SV-based nuclear tree

The input SV vcf file (homo_and_hetero_noDoublonsInCoordinates.vcf.gz) was generated using the non-redundant SV dataset. Based on the presence/absence information of these identified SVs in each assembly entry, a phylip-formatted 0/1 matrix was generated accordingly and used for tree building. IQtree (version 1.6.12; options: -s $prefix.phylip -st MORPH -m MK+ASC -bb 1000 -alrt 1000 -nt $threads - pre $prefix.iqtree -safe) was used to generate the phylogenetic tree. A thousand rounds of ultrafast bootstrap (UB) and approximate likelihood-ratio test (aLRT) were used to assess the branch supports.

#### Phylogenetic tree processing and visualization

For the phylogenetic trees generated above, tree-based operations such as rerooting, branch trimming, tip label extraction were performed by the nw_reroot, nw_prune, nw_labels tools from the Newick Utilities package (version: 1.6.0). Tree visualization was performed by the R package ggtree (version: 3.2.1). Cophylo comparison was conducted by the R package phytools (version: 1.0-3). The distance between trees was evaluated in terms of the quantity of information that the trees’ splits hold in common with The Clustering Information Distance implemented in the R package TreeDist (version: 2.4.1).

### Molecular dating

We used the formula for molecular dating published earlier (Fay and Benavides, 2005). We considered 100 and 365 generations per year to bound our estimations, as previously suggested (D’Angiolo *et al*., 2020). The value of the mutation rate of 2.31072123540072E-10 was calculated as the average of the rates for homozygous and heterozygous lines reported previously (Tattini *et al*., 2019). The pairwise distances between strains were calculated using MEGA11 (Tamura *et al*., 2021) as p-distance, using only the 4-fold degenerate sites. To determine if a codon position is a 4-fold degenerate site, we scan over every codon and codon position (i.e., 1st, 2nd, 3rd positions) based on NCBI’s codon table (https://www.ncbi.nlm.nih.gov/Taxonomy/Utils/wprintgc.cgi) based on the CDS alignment of each orthologous gene group. All codon positions corresponding to 4-fold degenerate sites were concatenated together to form the 4-fold degenerate site alignment of the corresponding CDS alignment. The 4-fold degenerate site alignment of all 1-to-1 ortholog CDSs were further concatenated to form a super alignment of 4-fold degenerate sites.

## LEGENDS TO SUPPLEMENTARY FIGURES

**Supp. Fig. 1:** Description of the 142 strains belonging to the ScRAP. The strains come from 3 datasets, (i) 100 de novo sequenced and assembled genomes, (ii) 18 re-assembled genomes using previously available raw Nanopore read data and (iii) 24 publically available complete genome assemblies, including the S288C reference genome. The number of strains with different ploidy and heterozygosity levels is indicated for each of the three datasets.

**Supp. Fig. 2:** Distribution of various assembly metrics. Each black dot corresponds to the value for one strain, the red dot shows the position of the S288C reference genome in the distribution. Median values are indicated by the gray crossbars. The horizontal dotted lines show the target values for complete assembly.

**Supp. Fig. 3:** Histogram of the percentage of sequence identity between each strain and the S288C reference genome, split by dataset.

**Supp. Fig. 4:** Comparison between the topologies of the ortholog-based (left) and SNP-based (right) phylogenetic trees.

**Supp. Fig. 5:** (A) Impact of haplotype phasing on SV validation for the different levels of ploidy. (B) Distribution of the number of heterozygous SV per strain split by ploidy. For diploids this was simply done by considering any variant that did not contain both HP1 and HP2 genomes as heterozygous. For polyploids, first the phased genomes were aligned to the reference, coverage was calculated around the region of the event and this coverage was then used to estimate the maximum number of haplotypes present. If the number of phased blocks validating the variant was fewer than the max haplotypes, the event was considered heterozygous.

**Supp. Fig. 6:** Proportions of SV calls, split by types, that contain repetitive (repeat) or unique (novel) sequences according to a BLAST search. The region involved in each event was searched for using BLAST within the reference genome and matches above an e-value of 0.001 and/or 90% base identity were removed. The remaining alignments needed to cover above 50% or 150% of the query length to be considered a repetitive region in insertions and deletion/duplication/inversions respectively. Less than 50% of the query coverage for an insertion would indicate that the majority of the region is not already present within the reference genome. Likewise, for a deletion/duplication/inversion, the region is already present in the reference genome by default and therefore 150% of the query would indicate that the majority of the region is also present elsewhere in the reference.

**Supp. Fig. 7:** Association between various genetic elements and SV breakpoints split by SV type. The breakpoints of all SVs within every ‘best’ genome were matched to the corresponding gff annotation file. These were then compared to the genome wide proportion of events to calculate a per genome enrichment. This was done by getting the breakpoint-associated features (left) and the closest element for intergenic breakpoints (right).

**Supp. Fig. 8:** (A) Number of the different types pf SVs present in the set of 51 isolates used to study the relationship between SVs and gene expression variation (Ins for insertions, Del for deletions, Inv for inversions, Dup for duplications, Transloc for translocations, and Contr for contractions). (B) Comparison of the expression level allowed to test for the impact of the SV. Left panel, comparison of the presence (+ SV) or absence (-SV) of a deletion event in the regulatory region of the ORF YHR043C. Right panel, comparison of the presence (+ SV) or absence (-SV) of a duplication of the ORF YHR054C.

**Supp. Fig. 9:** Comparison of the distributions of the aLRT node support values between the orthologue, SNP and SV-based phylogenetic trees.

**Supp. Fig. 10:** Accumulation of snifles (A) and paftools (B) derived SVs as a function of SNVs using.The categories ‘Heterozygous monosporic’ and ‘Homozygous monosporic’ correspond to monosporic isolates derived from the sporulation of heterozygous and homozygous parental diploid strains, respectively.

**Supp. Fig. 11:** Contribution of Non-Centromere Related (NCRs) coverage deviations in strains containing simple and complex aneuploidies. Strains with complex aneuploidies are more likely to have an NCR (left), are more likely to have a higher number of them (middle) and are more likely to have on average a larger (mean size) NCRs (right).

**Supp. Fig. 12:** Distribution of telomere length at all chromosome ends in each strain of the ScRAP split by dataset. The inset plot shows the distribution of the average telomere length per strain for each dataset.

**Supp. Fig. 13:** Telomere length properties. (A) scatter plot showing the positive correlation between the mean telomere length and its variance per strain. The Pearson correlation coefficient and its associated p-value are indicated. Boxplots showing the distribution of telomere lengths per strain split by zygosity (B), ploidy (C), and ecology (D) . Wilcoxon mean comparison p-values are indicated.

**Supp. Fig. 14:** Individual telomere sizes per chromosome in each of the 100 *de novo* sequenced and assembled strains from the ScRAP.

**Supp. Fig. 15:** Distribution of subtelomeric elements among the 100 *de novo* sequenced and assembled strains from the ScRAP. (A), (C) and (E) show the number of core X, Ty5 and Y’ elements found on each chromosome end across the 100 strains, respectively. (B), (D) and (F) show the distributions of the mean telomere length in the presence or absence of the corresponding subtelomeric elements across all (left) or TEL03L chromosome ends. Wilcoxon mean comparison p-values are indicated.

**Supp. Fig. 16:** Distribution of the Y’ elements across the ScRAP. (A) Histogram of the number of Y’ elements per chromosome end, split by dataset. (B) Histogram of the number of Y’ elements per strain, split by dataset.

**Supp. Fig. 17:** Genome annotation metrics. The panels corresponding to the diploid assemblies give the results separately for each haplotype (HP1 and HP2) while for the panels corresponding to the triploid and tetraploid assemblies, the results comprise annotations of all phased blocks together.

**Supp. Fig. 18:** Genome size variation across the ScRAP. Scatter plots showing the genome size of each strain, split by dataset, as a function of the number of (A) Y’ elements, (B) Ty elements and (C) Y’ + Ty elements.

**Supp. Fig. 19:** Distributions of the mean telomere length in the presence or absence of Y’ elements for each chromosome end across the 100 *de novo* sequenced and assembled strains from the ScRAP. Wilcoxon mean comparison p-values are indicated.

**Supp. Fig. 20:** The phylogenetic tree on the left is identical to Figure 1 and corresponds to a tree based on the concatenated protein sequence alignment of 1,618 1:1 ortholog. The green, red, blue and yellow symbols indicate the ecological origins. Ploidy levels and zygosity are symbolized by the shapes of the symbols as in Figure 1. The rectangles on the right indicate the distribution of the regions A to G across the 142 strains.

**Supp. Fig. 21:** (A) The HGT region B from *Zygosaccharomyces parabailii* was fully assembled in its native chromosomal position for the first time. Roughly 2 kb upstream the HGT (light grey boxes), we observed a 1 kb region (corresponding to YHR208W) with high local homology between *S. cerevisiae* and *Z. parabailii* (75% identical, overall), including several tracts between 10 and 20 bp of complete identity (green boxes). At the telomere side, this region shows a Y’ element, which we found also in the *Z. parabailii* assembly. The phylogenetic tree of the Y’ elements shows that the Y’ sequence in *Z. parabailii* is different from the one of the other *Saccharomyces* species and several copies of the Y’ BDM gave a close match, while other have an *S. cerevisiae*-type sequence. These data are consistent with a 3-way HGT exchange, with an initial transfer of Y’ (and perhaps associated sequences) from an undetermined species of the *Saccharomyces* genus to *Z. parabailii* followed by the transfer to *S. cerevisiae.* (B) Although the ORFs from region G, identified only in the ALI isolate, had been already described in Peter et al 2018, long-read assemblies enabled to describe the region as a genuine HGT and to identify the donor species. Protein sequence alignments reveal several ORFs with high sequence identity, up to 100% amino acid (AA) identity and 100% overlap (i.e. ORF 0042950) with *L. fermentati* (Lf). Lower identity values are observed also in the alignments against *L. dasiensis* (Ld). (C) Region F from an unknown source was retrieved in the assemblies of two strains. Remarkably, it is found both in the canonical subtelomeric position (e.g. in AEG) as well as reduced incore-chromosomal positions (in the CPA isolate).

**Supp. Fig. 22:** Distribution of the tRNA gene families in the ScRAP. (A) Number of tRNA genes annotated in genomes split by publication. The names of the first authors of the following publications are indicated (Bendixsen *et al*., 2021; Berlin *et al*., 2015; Czaja *et al*., 2020; Goffeau *et al*., 1996; Jenjaroenpun *et al*., 2018; Shao *et al*., 2018; Yue *et al*., 2017; Zhang and Emerson, 2019). (B) Number of tRNA annotated in homozygous/haploid genomes (no) and collapsed assemblies from heterozygous genomes (yes). (C) Conservation of tRNA gene families across 100 isolates. (D) Relative chromosomal location of conserved, gained and lost tDNA gene families.

**Supp. Fig. 23:** (A) Number of TE sequences per strain across the 142 haploid/collapsed genome assemblies. All sequences from the 5 Ty families are pooled together by category. (B) Number of complete Ty elements per strain across the 142 haploid/collapsed genome assemblies. (C) Distribution of the 126 insertion sites across the 100 haploid or homozygous genomes considering either the complete Ty elements or all types of TE sequences (complete, truncated and solo-LTRs). (D) Scatter plot between number of solo-LTRs per insertion site and the number of strains sharing an insertion site.

**Supp. Fig. 24:** Map of the *de novo* insertions of complete Ty elements across the 100 homozygous explored strains. The map shows the 61 insertion sites in which only complete elements are found and never soloLTRs, which strongly suggests that these sites correspond to recent insertions. Strains are organized according to the nuclear phylogenetic tree (Figure 1).

**Supp. Fig. 25:** Schematic representation of all processes applied to the ScRAP.

**Supp. Fig. 26:** Comparison of using the reference versus the de-novo assemblies using three statistics, the number of heterozygous variants that were phased, the nV90 and the number of phased blocks. All 3 metrics move in the direction expected for better phasing quality.

**Supp. Fig. 27:** Validation of SV calls. Validation requires a call to be found within at least two genomes, whether within or between strains. The curves show the evolution of the proportion of validated calls as a function of the window size used to consider that two breakpoints were overlapping. The categories are as follows, the SV is only validated by genomes from the same strain (within single strain). The SV is validated by being found in multiple genomes for 1 or more strains, plus being found in multiple strains (within and across strains) and the SV is validated across strains without having multiple genomes for any of those strains (only across strains).

**Supp. Fig. 28:** Proportion of validated SV split by publication. The names of the first authors of the following publications are indicated (Bendixsen *et al*., 2021; Berlin *et al*., 2015; Czaja *et al*., 2020; Goffeau *et al*., 1996; Jenjaroenpun *et al*., 2018; Shao *et al*., 2018; Yue *et al*., 2017; Zhang and Emerson, 2019).

## LIST OF SUPPLEMENTARY TABLES

**Supp_Table1_nuclear_genome_stats.xlsx**

**Supp_Table2_strain_origin.xlsx**

**Supp_Table3_mito_genome_stats.xlsx**

**Supp_Table4_molecular_clock.xlsx**

**Supp_Table5_SVs_final_noseq.xlsx**

**Supp_Table6_SNP_and_SV_per_strain.xlsx**

**Supp. Table 7 SV_Expression**

**Supp_Table8_aneuploidy.xlsx**

**Supp_Table9_aneuploidy_large.xlsx**

**Supp_Table10_aneuploidy_large_NCR.xlsx**

**Supp_Table11_annotation_stats.xlsx**

**Supp_Table12_HGT.xlsx**

**Supp_Table13_tRNA_100_strains.xlsx**

**Supp_Table14_tRNA_anticodon.xlsx**

**Supp_Table15_Ty_insertion_sites.xlsx**

**Supp_Table16_sequencing_stats.xlsx**

**Supp_Table17_subtelomeres.xlsx**

